# Inferring differentiation trajectories from T cell clonotype distributions

**DOI:** 10.64898/2026.06.29.735205

**Authors:** Mariia Guryleva, Maximilian Danielsson, Christopher Andrew Tibbitt, Jonathan Coquet, Ben Murrell

## Abstract

Single-cell RNA sequencing (scRNA-seq) enables reconstruction of cellular differentiation trajectories, but most trajectory inference methods rely largely on gene expression and overlook lineage relationships. In T lymphocytes, the T cell receptor (TCR) provides a unique, endogenous barcode that is stably inherited during clonal expansion. Here we introduce PhyloTrajectory, a framework that leverages TCR-based clonotype frequencies together with scRNA-seq to infer T cell differentiation dynamics. By modeling clonotype frequency evolution as a continuous stochastic process, PhyloTrajectory reconstructs differentiation tree topologies from clonotype frequencies across cellular subsets. We investigate the ability to recover the underlying tree topology using simulated data, and apply our approach to CD4^+^ T cells in murine models of allergic inflammation and viral infection. We further demonstrate that PhyloTrajectory is useful in the context of exogenously introduced barcodes. Overall, this shows that a stochastic model of clonal expansion can be used to infer cell state transitions from clonotype frequencies.

## Introduction

Single-cell technologies have revolutionized our ability to characterize cell subsets and individual cell states [1]. While the widely used single-cell RNA sequencing (scRNA-seq) provides detailed information about the transcriptomic profiles of cells, it can also be complemented with data on chromatin accessibility, surface proteins, and adaptive immune repertoires [2], [3], [4]. These multimodal features precisely characterize cell phenotypes and enable computational study of cellular dynamics. Over 70 trajectory inference methods have been developed to reconstruct differentiation paths, with Monocle, Slingshot, and PAGA being among the most widely used [5], [6]. Despite multimodal data availability, most trajectory inference methods rely solely on gene expression [7].

However, gene expression states of progenitors do not always predict their eventual fates. In such cases, lineage tracing provides a more direct view of differentiation [8]. DNA barcodes present in progenitor cells are inherited by descendant cells, allowing for the determination of individual differentiation lineages. While these techniques can recover actual differentiation lineages, they are technically demanding and subject to several limitations [8]. First, these methods are labor-intensive. Additionally, they are affected by stochastic barcode insertion effects and typically limited to small numbers of model organisms [8], [9].

T cells are central to adaptive immunity, mediating responses to infections, allergens, and neo-antigens in the context of cancer. Through genetic recombination of the T cell receptor (TCR) locus and the addition of non-templated nucleotides at the CDR3 region of the TCR, highly diverse T cells develop from the thymus and seed the periphery [10]. Upon encountering cognate antigen, naive T cells undergo clonal expansion and differentiation into effector or memory states [11], [12]. Crucially, unlike B cell receptors, the TCR sequence remains unchanged throughout the lifetime of a T cell lineage. The diversity of TCR sequences enables them to serve as endogenous barcodes for clonally related cells, eliminating the need for exogenous constructs.

Recent computational approaches have begun to leverage clonal identity to study differentiation beyond gene expression. For example, clone2vec captures continuous variation in clonal fate biases by embedding clones based on the transcriptional similarity of their constituent cells [13]. However, since many differentiation patterns are inherently tree-like, phylogenetic approaches offer a natural alternative framework. Classical phylogenetic methods use probabilistic models of sequence evolution to infer relationships by finding trees that best explain observed sequences [14], [15]. While TCR sequences remain fixed during clonal expansion, clonotype *frequencies* change over time, and serve as footprints of antigen exposure and cell differentiation [16], [17].

Here we present PhyloTrajectory, a statistical framework that infers differentiation tree topology and branch lengths by exploiting clonotype frequency distributions across transcriptional states. We benchmark PhyloTrajectory on simulated data and apply it to CD4^+^ T cells in murine allergy and viral infection models, revealing branching patterns in T helper differentiation. Finally, we demonstrate application of our model to lineage-barcoded data, showing its generality beyond TCR-based systems.

## Results

### Overview of the PhyloTrajectory framework

Classical phylogenetic models infer evolutionary relationships from sequence mutations, where the pattern of inherited mutations contains sufficient information to allow us to infer the evolutionary tree (Figure 1, c left panel). In our setting, while TCR sequences remain static, clonotype frequencies change dynamically as cells proliferate and differentiate. If a clonal expansion event occurs prior to differentiation, then all else being equal the expanded clonotype is more likely to occur at higher frequencies in the post-differentiation descendant cellular states (Figure 1, a). Thus the clonotype frequency changes are heritable over the differentiation tree and, just like sequence mutations in phylogenetics, the distribution induced by this inheritance leaves an imprint of the tree upon the observed clonotype frequencies. We exploit this to reconstruct the differentiation tree itself (Figure 1, b).

**Figure 1:**
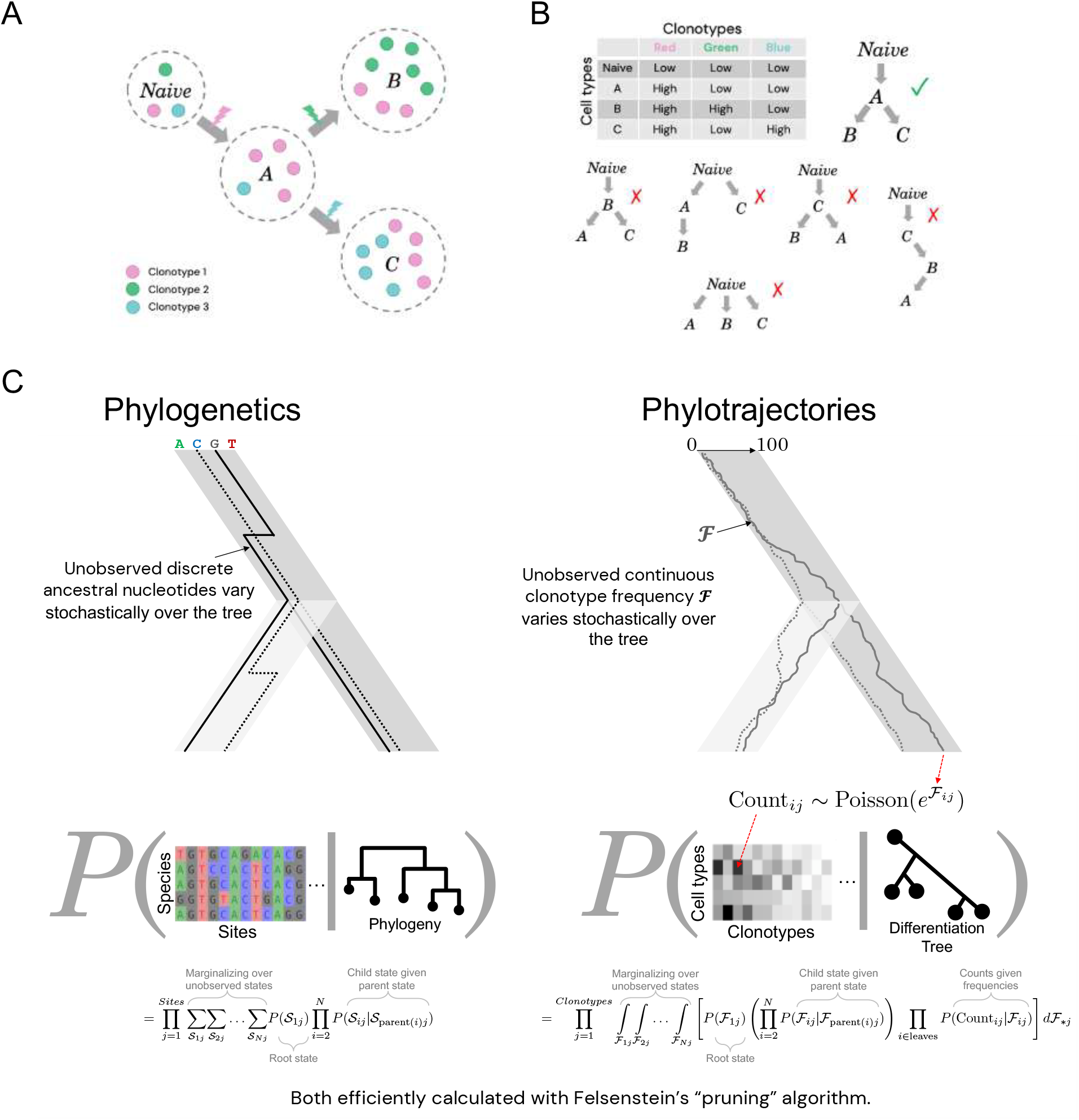
The PhyloTrajectory framework. **a.** Schematic of T cell differentiation following antigen encounter. Naïve T cells undergo clonal expansion and differentiate into distinct effector states (A, B, C); three representative clonotypes are shown. **b**. Illustration of the relationship between clonotype count matrices and candidate differentiation trees. For a given count matrix (rows: cell types; columns: clonotypes), only the tree that correctly groups cell states sharing clonotype abundances is compatible with the differentiation process (A). **c**. Parallel between classical phylogenetic inference (left) and PhyloTrajectory (right). In phylogenetics, unobserved discrete ancestral nucleotide states evolve stochastically over the tree; in PhyloTrajectory, unobserved continuous clonotype log-frequencies evolve over the differentiation tree. Both likelihoods marginalise over unobserved internal states and are computed efficiently using Felsenstein’s pruning algorithm.

PhyloTrajectory models the unobserved latent clonotype frequency as a continuous stochastic process evolving over a differentiation tree. Observed cell-state clusters correspond to the leaves of the tree, whereas internal nodes represent unobserved, ancestral states from which differentiation bifurcations give rise to descendant states. Branch lengths quantify the expected amount of clonotype-frequency divergence between connected states under the model, rather than chronological time. At the leaves, the unobserved latent clonotype frequencies are linked to the observed clonotype counts through a sampling distribution.

Our observed data are a matrix of clonotype counts (ie. non-negative integers), with one value per cell state per clonotype. The differentiation tree, not directly observed, is a hypothesis about the relationships between cell states.

As described in Methods, we model the logarithm of the latent clonotype frequencies (by which throughout we mean abundances, not proportions), allowing stochastic exponential growth or decay, with an Ornstein-Uhlenbeck (OU) process [18]. This is a simple continuous diffusion process with an equilibrium distribution, which is needed to prevent the overgrowth of clonotypes with time under the model. This process is described by the mean (log) frequency of a random clonotype, the mean-reversion rate, and the process volatility. We allow uncertainty in all of these parameters, and the tree itself, as well as the frequency distribution at the root of the tree. At the leaves of the tree, corresponding to extant cell states, for an underlying frequency the count is approximately Poisson distributed due to sampling a random subset of cells.

Given a candidate tree topology and branch lengths, together with a specification of the continuous stochastic process that governs clonotype frequency dynamics, we can adapt a classic algorithm from phylogenetics [14] to efficiently compute the likelihood of an observed matrix of clonotype counts (see Methods; Figure 1, c right panel). With this efficient likelihood calculation, we can use standard tools from Bayesian inference to sample from the posterior distribution of trees and process parameters, which characterizes the space of differentiation trajectories that are compatible with the observed data.

### Simulations

With any new statistical model, there is a question of whether, and to what extent, the data can inform us of the quantity we are interested in. Here we use idealized simulations, both for realistic sample sizes and in the large-sample regime, to investigate whether phylogenetic relationships can be reconstructed from clonotype count matrices alone. Synthetic datasets were generated by simulating clonotype counts along predefined tree topologies with between 4 and 10 leaves.

For a fixed branching structure, clonotype frequencies were propagated along the tree under a Brownian-motion process, and count matrices were obtained by multinomial sampling at the tips, both representing mild violations (Brownian vs OU, and Multinomial vs Poisson) of the specific modeling choices used during reconstruction by PhyloTrajectory (Figure 2, a). The simulated count matrices were then used as input for the PhyloTrajectory model to recover the tree structure, comparing the inferred topologies against ground truth.

**Figure 2:**
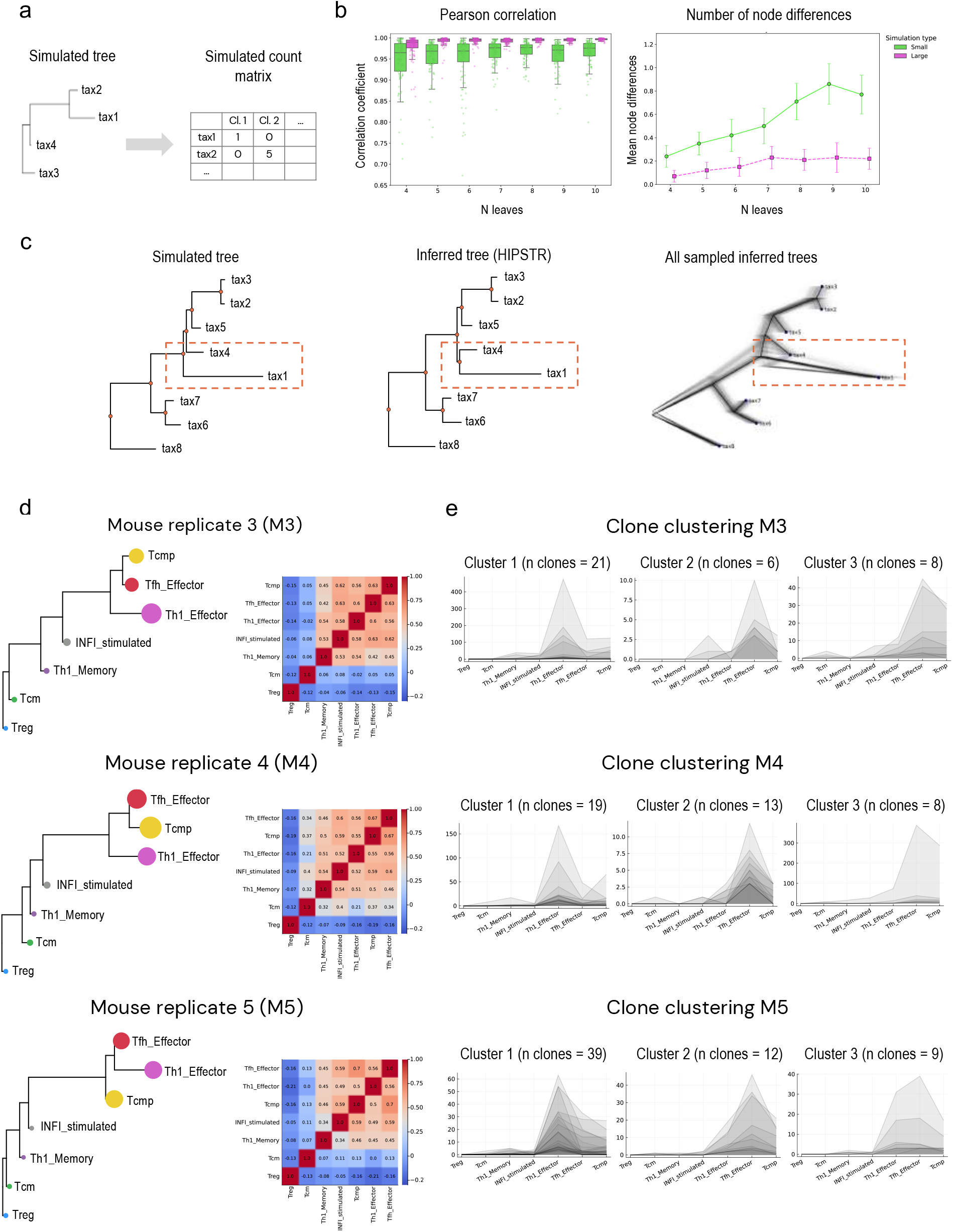
Simulation benchmarking and application to LCMV infection. **a.** Schematic of the simulation framework. Clonotype frequencies were propagated along a known tree topology by Brownian motion and converted into tip-level clonotype count matrices by multinomial sampling. **b**. Reconstruction accuracy across 100 simulated datasets per tree size, using trees with 4–10 leaves and two simulation regimes. Pearson correlation between inferred and simulated pairwise branch-length distances (left); boxplots summarize the distribution across replicates, with individual simulations shown as points. Topological reconstruction error, measured as the number of node differences between inferred and simulated trees (right); points show the mean across replicates and error bars indicate 95% confidence intervals. Colours indicate the number of leaves, while hatching or line style denotes the simulation regime. **c**. Example of a partially unresolved inference. The simulated tree, shown on the left, and the highest posterior summary tree (HIPSTR) consensus tree, shown in the centre, differ at a single internal node, highlighted by the dashed box. Posterior uncertainty at this node is reflected by the spread of Markov chain Monte Carlo(MCMC)-sampled trees shown on the right. **d**. PhyloTrajectory-inferred differentiation trees and pairwise Pearson correlation heatmaps of log-transformed clonotype counts across cell states for three biological replicates, M3, M4 and M5, from a lymphocytic choriomeningitis virus (LCMV) infection model. **e**. Selected clonal clusters for each replicate, showing observed clonotype counts across cell states. The number of clonotypes assigned to each cluster is indicated above each panel.

We ran two classes of simulations: a realistic sample size setting (mean 15,000 cells), and a large sample regime (mean ∼ 1 million cells). Reconstruction accuracy was quantified with two complementary metrics: (i) the Pearson correlation between the vectors of all pairwise branch-length distances and root-to-leaf distances (to make the metric sensitive to root placement inference) of the inferred and simulated trees, and (ii) the number of unmatched internal nodes (“Node Diff”), normalized to represent unique topological splits.

Across 100 replicate simulations per leaf count, the inferred trees closely reproduced the ground-truth topologies in both settings. Branch-length distances were strongly correlated with the truth, with a mean Pearson correlation of 0.96 in the small setting and above 0.99 in the large setting (Figure 2, b; Supplementary Table S1). Topological recovery was likewise high: the mean number of node differences was 0.55 (small) and 0.18 (large), more than one topological discrepancy occurred in only 11.1% of small-setting and 1.9% of large-setting replicates (Figure 2, b; Supplementary Table S1). Accuracy depended on the number of leaves (representing number of cell states), and this dependence was more pronounced in the small than in the large setting (Figure 2, b).

The few topological mismatches were concentrated in subtrees with very short internal branches. In these regions the likelihood is nearly invariant between two competing representations of the same event: (i) a leaf that corresponds to an internal state (no split), or (ii) a rapid bifurcation followed by almost no evolution along one descendant branch. When the internal edge is shorter than the scale of the observational noise, the data provide insufficient signal to distinguish these cases, and the MCMC posterior assigns non-negligible probability to both (Figure 2, c). This identifiability limit explains the isolated node discrepancies, while branch-length correlations remain high overall in these samples.

Together, these results indicate that, under the tested conditions, PhyloTrajectory recovers tree topologies with high accuracy and branch lengths that correlate strongly with the ground truth.

### Consistent differentiation trajectories across LCMV replicates

We next applied the PhyloTrajectory inference framework to TCR repertoires from an acute LCMV (Armstrong) infection model, profiled at the peak (day 10 post-infection) of the immune response. This dataset comprises virus-specific GP66-77 CD4^+^ T cells isolated from spleens of five infected mice that underwent identical experimental treatment and sequencing protocols [19].

The original *Khatun et al*. dataset was re-processed and re-clustered prior to analysis. We annotated cell states with ProjecTILs by projecting the cells onto the published reference atlas of virus-specific CD4^+^ T cells spanning acute and chronic infection [20]. Projection recovered the nine reference cell states defined in the atlas, represented across all five mice: Th1 effector (Th1_Effector), Th1 memory (Th1_Memory), Tfh effector (Tfh_Effector), Tfh memory (Tfh_Memory), central memory (Tcm), central memory precursors (Tcmp), regulatory T cells (Treg), IFN-stimulated Tfh-like cells (INFI_stimulated), and an Eomes-high (Eomes_HI) population. The projected annotations recapitulated the canonical marker patterns reported for these states (Supplementary Figure S1, a). Consistent with prior characterizations, Th1 effector cells showed elevated expression of *Cxcr6* and *Ly6c2*, whereas Tfh effector cells preferentially expressed *Cxcr5* and *Slamf6*. Tcmp cells were enriched for *Ccr7*, consistent with a central memory precursor phenotype. Memory populations (Th1_Memory, Tfh_Memory, and Tcm) exhibited higher expression of *Tcf7* and *Il7r*, in agreement with their memory-associated transcriptional program. Regulatory T cells expressed *Foxp3* and associated markers, while the INFI_stimulated cluster was characterized by elevated interferon-stimulated genes. The Eomes_HI population showed increased expression of *Eomes* and cytotoxic-associated genes such as *Gzmk*.

Overall, mice M1 and M2 contained lower total numbers of T cells and displayed a relative enrichment of memory subsets (Th1_Memory and Tfh_Memory) compared to the other replicates. In contrast, mice M3–M5 exhibited higher total cell counts and more comparable distributions of T cell populations (Supplementary Figure S1, b). We thus focused subsequent trajectory inference on M3, M4, and M5. These three replicates also showed highly comparable clonal diversity, with similar numbers of unique clonotypes (M3: 698; M4: 747; M5: 718); the distribution of expanded clonotypes across cell states is shown in Supplementary Figure S1, c.

The differentiation trees inferred using PhyloTrajectory captured a mostly-consistent global differentiation structure across the three analysed replicates. In all mice, Treg and Tcm occupied root-proximal positions, whereas Th1_Memory and IFN-stimulated cells were placed along the internal backbone upstream of the terminal effector-associated states, and Th1_Effector, Tfh_Effector, and Tcmp occupied distal branches. The pairwise clone-correlation heatmaps in Figure 2, d were consistent with this structure, with highly correlated cell states generally placed on neighbouring branches and root-proximal states showing weaker correlations with the distal effector-associated populations; the corresponding confidence-colored inferred trees are shown in Supplementary Figure S1, d.

The IFN-stimulated cell state was consistently positioned along the internal backbone, prior to the split among distal effector-associated states. This placement is compatible with a transitional state whose clonotype composition has not diverged toward a single terminal fate. Treg was consistently the most root-proximal leaf, consistent with limited divergent clonal expansion and weak clonal association with distal effector states. Tcm also occupied a root-proximal position, but was generally placed downstream of Treg, due to broader clonal sharing with other non-terminal states.

The specific terminal arrangement of Th1_Effector, Tfh_Effector, and Tcmp was consistent between M3 and M4, where Tfh_Effector and Tcmp coalesce first, but the branching orders is swapped in M5, with Tfh_Effector and Th1_Effector coalescing first. However, the Bayesian posterior credibility of the Tfh_Effector and Th1_Effector coalescence is only 77% (see Supplementary Figure S1, d), so the model is not asserting that there is statistical evidence that M5 differs from M3 and M4 here.

To examine clonotype-level patterns, we applied a clustering algorithm (Dirichlet Process-means [21], [22]) to the latent frequency distributions inferred at the nodes and leaves of the posterior consensus tree, conditioned on the observed counts (see Methods). This shows that the tree similarity is driven by a common pattern of clonal expansion, where in each replicate, selected clone clusters were enriched in distal effector-associated states, including Th1_Effector-biased and Tfh_Effector-biased clones. These patterns are consistent with the distal placement of Th1_Effector and Tfh_Effector in all three inferred trees.

The relationship between Tcmp and the effector-associated states differed between replicates, but with low statistical confidence. In M3 and M4, selected clone clusters showed enrichment across Tfh_Effector and Tcmp, consistent with the inferred grouping of these two states in the corresponding trees. In M5, Tcmp remained closely associated with the distal effector region, but was placed on a short branch upstream of the Th1_Effector–Tfh_Effector pair. However, this pattern appears to be driven by a small number of clones expanded in both Th1_Effector and Tfh_Effector states, with slightly less representation in Tcmp. This is consistent with the low confidence associated with this difference between M5 and M3/M4.

Overall, across three biological replicates, PhyloTrajectory inferred a consistent global differentiation organization, with Treg and Tcm positioned near the root, IFN-stimulated cells along the internal backbone, and Th1_Effector, Tfh_Effector, and Tcmp occupying distal branches. The precise terminal arrangement was consistent between M3 and M4, but different slightly in M5, although without statistical confidence in this difference.

### Differentiation trajectories in allergen-driven CD4^+^ T cell responses

We generated a House Dust Mite (HDM) model dataset, where CD3+CD4+ T cells were isolated from bronchoalveolar lavage (BAL) on day 15. T cells were profiled with single-cell GEX and VDJ 10x pipelines. For comparison we also used a previously published Dog Allergen (DA) dataset (GSE244722) with a matching experimental setup, where murine BAL CD3+CD4+ T cells were profiled after administration of dog allergen extracts.

Unlike the LCMV model, where gp66-77 tetramer+ cells were purified and analysed, both allergen datasets comprise unenriched total CD4^+^ T cells. This demonstrates that PhyloTrajectory can be applied to an active immune response without antigen-specific cell isolation, so long as sufficient clonal expansion is present to sample detectable frequency covariation across states.

Both HDM and DA datasets were processed separately using uniform scRNA-seq/scTCR-seq workflows, and clonotype-resolved T helper (Th) populations were identified as described in Methods. The Dog Allergen model yielded eleven transcriptionally distinct CD4^+^ subsets, including Th2, Th17, multiple activated (Act_Tcf1+, Act2 and Act3) states, T follicular helper (Tfh), Naïve, T regulatory (Tregs), Th effector CTL-like (Th_CTL_like), IFN-response, and proliferating clusters comprising 1,719 unique clonotypes in total (Figure 3, d; Supplementary Figure S2). In contrast, the HDM model exhibited seven major clusters with primarily Th2-driven responses (Th2, Th2_2 clusters), Tregs, activated state (Act_Tcf1+), IFN-response, Naïve, and Proliferating populations spanning 2,017 unique clonotypes (Figure 3, a; Supplementary Figure S3).

**Figure 3:**
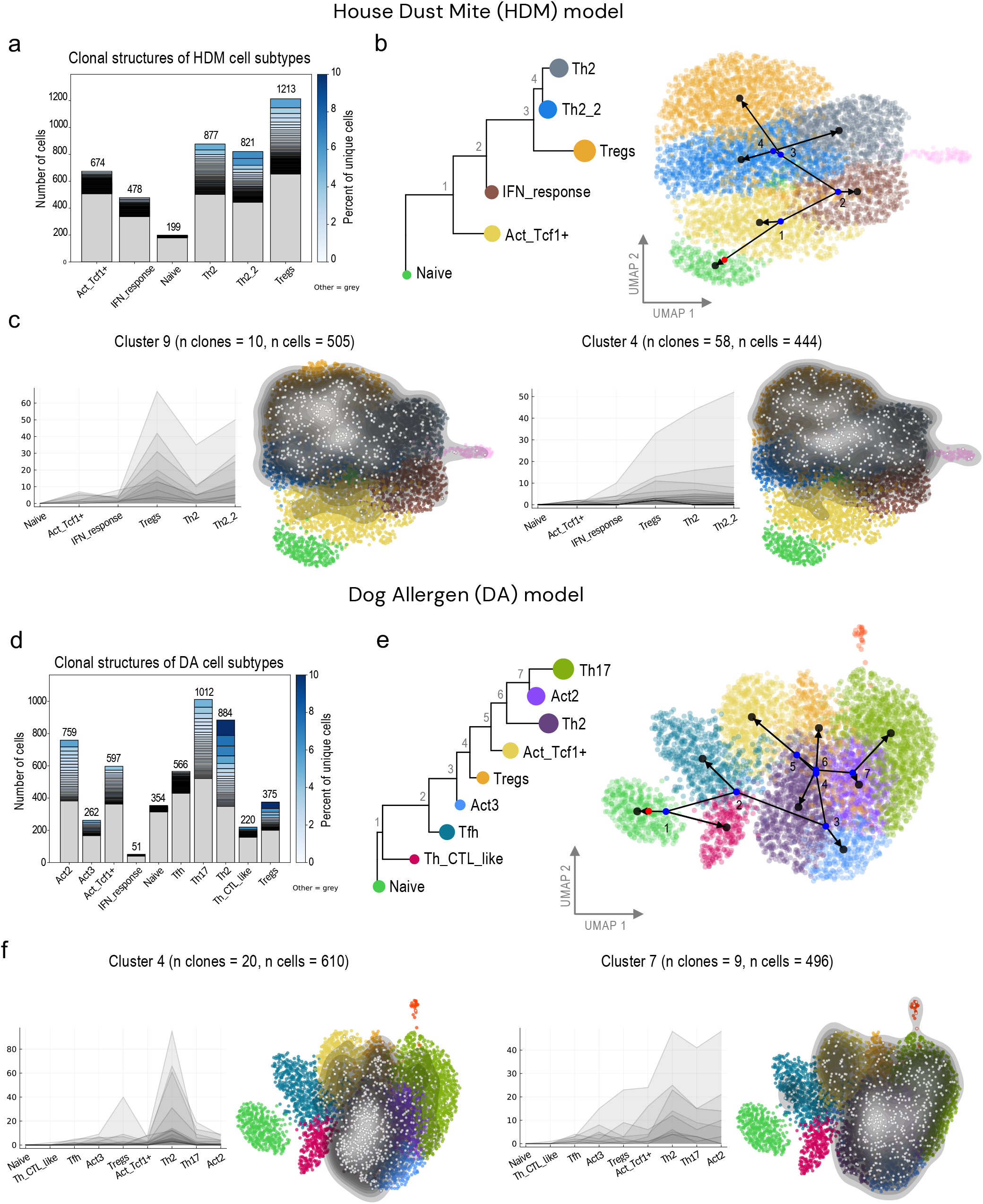
PhyloTrajectory inference applied to allergen-driven CD4^+^ T helper cells in house dust mite and dog allergen models. **a.** Clonal structures of HDM cell subtypes (excluding the Proliferative cluster). The bar plot shows the total number of cells per cluster, with stacked bars indicating the top 10% most expanded clonotypes; color intensity reflects the percentage of cells contributed by each clonotype. **b**. Differentiation topology inferred by PhyloTrajectory for the HDM model, shown as a consensus tree (left) and with the inferred trajectory overlaid on the UMAP embedding (right). **c**. Representative clonal clusters identified in the HDM dataset. For each cluster, the line plot (left) shows observed cell counts across cell types for individual clonotypes, and the density contour overlay (right) shows the spatial distribution of these cells in UMAP space. The number of clonotypes (n clones) and total cells (n cells) per cluster is indicated above each panel. **d–f**. As in **a–c**, for the dog allergen (DA) model.

The trajectories obtained with PhyloTrajectory reconstruction showed overall consistent tree structures between the two allergen models (Figure 3, b, e). Naïve populations were placed at the root and effector populations (Th2 and/or Th17) located at the distal branches. The inferred trajectories recapitulated expected differentiation hierarchies, but notable divergences were observed between the disease models. Specifically, the Treg populations occupied distinct positions in the differentiation trees: in HDM, Tregs emerged adjacent to effector Th2 populations, forming a relatively long separate branch indicative of shared clonotypes with the dominant Th2 axis. The long Treg branch indicates sub-stantial clonal divergence from the adjacent Th2/Th2_2 branch despite detectable clone sharings. Conversely, in the Dog Allergen model, Tregs appeared closer to the tree root with shorter branch length. This suggests that Tregs in this model, while diversifying earlier from effector states, did not include substantial clonal differences. These topological contrasts were corroborated by pairwise Pearson correlations of clonal compositions, confirming the agreement between clonal similarity matrices and phylogenetic distances (Supplementary Figure S4, a and b).

In HDM, Cluster 9 (n clones = 10, n cells = 505) and Cluster 4 (n clones = 58, n cells = 444) showed patterns of expansion across Treg and Th2/Th2_2 populations (Figure 3, c); we also identified a Treg-enriched cluster, Cluster 3 (n clones = 40, n cells = 343), shown in Supplementary Figure S4, c. This Treg-enriched HDM pattern was not mirrored by a Treg-only cluster in the displayed Dog Allergen clonal clusters. In Dog Allergen, Cluster 7 (n clones = 9, n cells = 496) spanned activated, effector, and Treg populations, including Act_Tcf1+, Th2, Th17, Act2, and Tregs, while largely excluding Th_CTL_like, Tfh, and Naïve. Cluster 4 (n clones = 20, n cells = 610) showed enrichment across Tregs and Act_Tcf1+ with lower representation in effector states (Figure 3, f).

In summary, both allergen models produced consistent global tree structures with Naïve at the root and effector populations at terminal branches, but differed in Treg positioning and clonal dynamic patterns between Treg and effector populations.

### Clonal heterogeneity within the HDM Treg compartment

The branch leading to the Treg cluster appeared unusually long relative to other leaves. A similar pattern is observed in phylogenetics when a recombinant sequence, with multiple distinct inheritance patterns for different regions of its genome, is placed on a single tree. In our setting, the analog of recombination would be where cells within one descendant cellular cluster arise from multiple distinct parent states, which can occur when cells are clustered too coarsely. When this occurs, one possible signal is that, within the descendant cluster, there is some association between clonality and expression state, where regions within the cluster exhibit different degrees of clonal expansion compared to other regions. We constructed a permutation test that aims to identify clusters where clonality varies, within a single cluster, across PCA space (see Materials and Methods).

Among all clusters in the HDM dataset, the Treg cluster showed markedly higher signal for such clonal heterogeneity, indicating the presence of distinct subpopulations (Figure 4, a). In contrast, no cluster in the Dog Allergen dataset exceeded the permutation test’s null distribution (Supplementary Figure S5, b; Supplementary Table S2).

**Figure 4:**
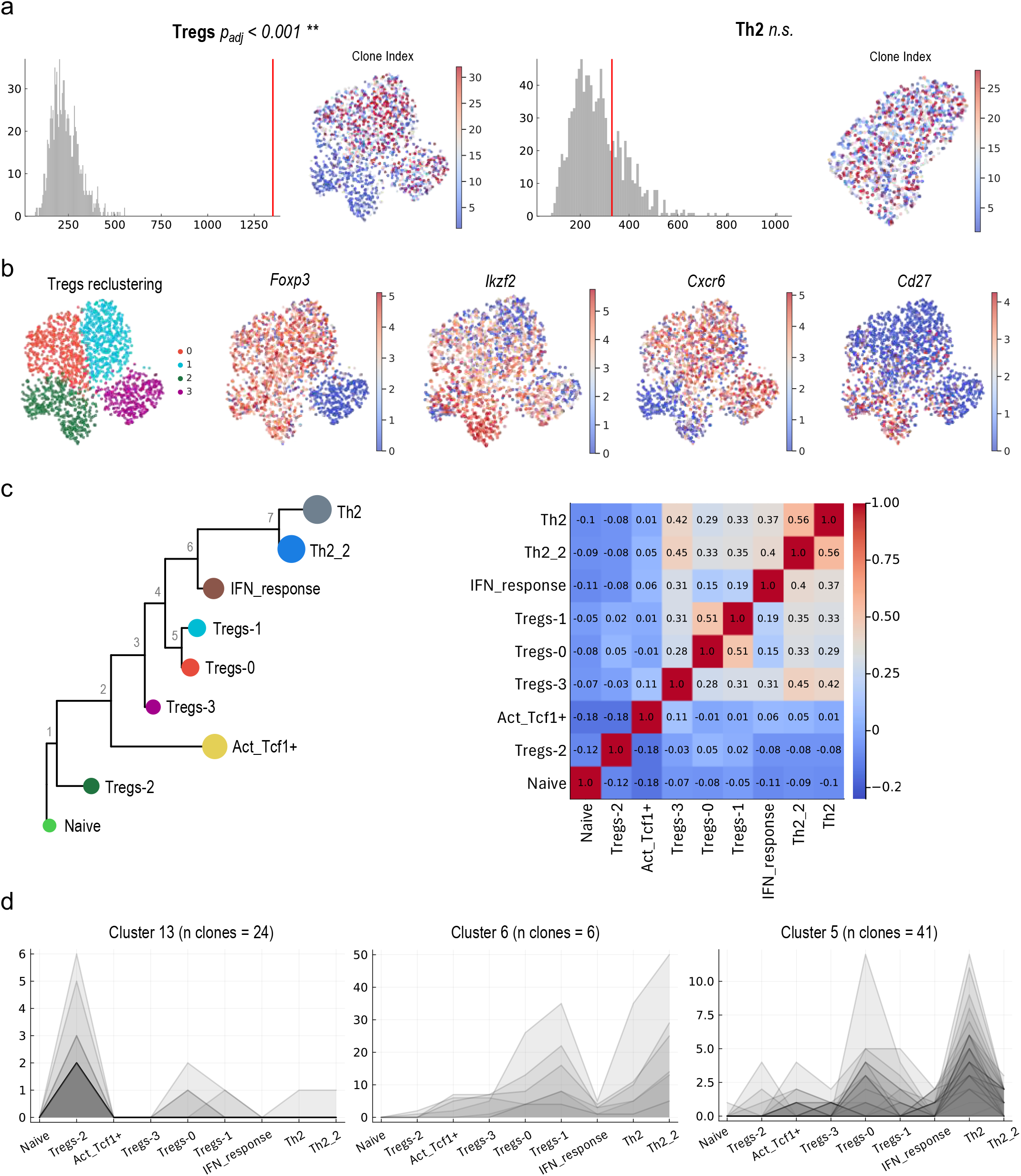
Clonal heterogeneity within the HDM Treg compartment. **a.** Representative nearest-neighbour permutation tests for local clonal size heterogeneity in HDM transcriptional clusters. For each cluster, the observed variance of the local-neighbourhood maximum clonotype size in PCA space is shown by the red vertical line and compared with the null distribution generated from 1,000 within-cluster clone-label permutations, shown as grey histograms. Adjacent Clone Index maps show the clone size associated with each cell, defined as the size of the clonotype to which that cell belongs. Empirical one-sided permutation p-values were adjusted across clusters using the Bonfer-roni-Holm procedure. Tregs showed significantly elevated local clonal size heterogeneity, whereas Th2 cells did not. *p*_adj_, Bonferroni-Holm adjusted permutation p-value; n.s., not significant; \**p*_adj_ < 0.05; \*\**p*_adj_ < 0.01. **b**. UMAP of the HDM Treg population after re-clustering. The left panel shows the four Treg subpopulations, and the remaining panels show expression of selected Treg-associated marker genes: *Foxp3, Ikzf2, Cxcr6*, and *Cd27*. **c**. PhyloTrajectory-inferred differentiation tree after resolving the original Treg population into four Treg subpopulations (left), together with the corresponding pairwise Pearson correlation matrix of log-transformed clonotype counts across cell states (right). **d**. Selected clonal clusters showing observed clonotype counts across cell states. Grey profiles represent individual clonotypes within each cluster, and the number of clonotypes assigned to each cluster is indicated above each panel.

We therefore re-clustered the Treg population into four distinct subgroups: Tregs-0, Tregs-1, Tregs-2, and Tregs-3 (Figure 4, b). Following re-clustering, none of the four Treg subpopulations showed significant clonal heterogeneity by the permutation test (Supplementary Figure S5, b). Tregs-0 and Tregs-1 were both *Foxp3*+ with similar baseline expression patterns. Tregs-0 additionally expressed *Isg15, Ifit1*, and *Gzmb*, suggesting an activated or interferon-exposed state. Interferon-stimulated gene expression and granzyme B production have been associated with effector Treg function, including cytotoxic suppressive mechanisms in inflammatory and tumor microenvironments [23], [24], [25], [26]. Tregs-2 expressed *Foxp3* alongside elevated *Ikzf2* (*Helios*) and *Cd27*. Helios has been described as a marker of thymic-derived Tregs and is associated with enhanced *Foxp3* stability and suppressive function [27], [28]. *Cd27* expression further suggests memory-like properties and functional stability [29]. This subset may represent thymic-derived and highly stable Tregs. Cells in Tregs-3 were *Foxp3*-low but enriched for expression of *Maml2, Ccr5*, and the long non-coding RNA *Dleu2. Ccr5* is a chemokine receptor associated with lung homing and retention at inflammatory sites [30], [31]. *Dleu2* has been implicated in iTreg differentiation [32]. The low level of Foxp3 in this cluster suggests these cells may represent either destabilized “ex-Tregs” that have lost *Foxp3* expression, or precursor cells that have not yet acquired stable *Foxp3* expression during peripheral induction.

We further ran PhyloTrajectory with newly defined clusters. Interestingly, Tregs-2, the subset expressing *Helios*, was placed close to the root while the rest of Treg clusters were placed further down on the tree (Figure 4, c). Tregs-2 showed almost no clonal expansion; we identified a small group of clones specific to this population (Figure 4, d). Tregs-0, Tregs-1, and Tregs-3 lacked uniquely specific clones and mostly shared clonotypes with effector populations (Th2 and Th2_2). These clusters also exhibited higher levels of clonal expansions. Additional clonal clusters and trajectory overlays after Treg re-clustering are shown in Supplementary Figure S6.

Together, these analyses show that the initial Treg compartment in the HDM model was not a single homogeneous leaf on the phylotrajectory, but instead comprised clonally and transcriptionally distinct subpopulations which occupy distinct niches on the inferred trajectory.

### Application to LARRY lineage barcoding data

We applied our PhyloTrajectory framework to the *in vitro* LARRY (Lineage And RNA RecoverY) barcoding dataset from Weinreb et al. (2020) [33]. The dataset contained previously annotated hematopoietic lineages, including erythroid (Eryth), megakaryocyte (Mk), basophil (Baso), mast cell (Mast), eosinophil (Eos), neutrophil (Neu), monocyte (Mono), plasmacytoid dendritic cell (pDC), Ccr7^+^ migratory dendritic cells (Ccr7_DC), and lymphoid precursors (Lymph) (Supplementary Figure S7).

Since cell counts were large, we used a fine-grained clustering strategy to avoid heterogenous groups, and we inferred phylotrajectories independently for wells derived from two starting progenitor populations — multipotent Lin^−^Sca1^+^Kit^+^ (LSK) and oligopotent Lin^−^Sca1^−^Kit^+^ (LK). Across biological replicates, the reconstructed lineage topologies were broadly consistent, supporting robustness of the method to sampling variability(Figure 5; Supplementary Figures S8, S9).

**Figure 5:**
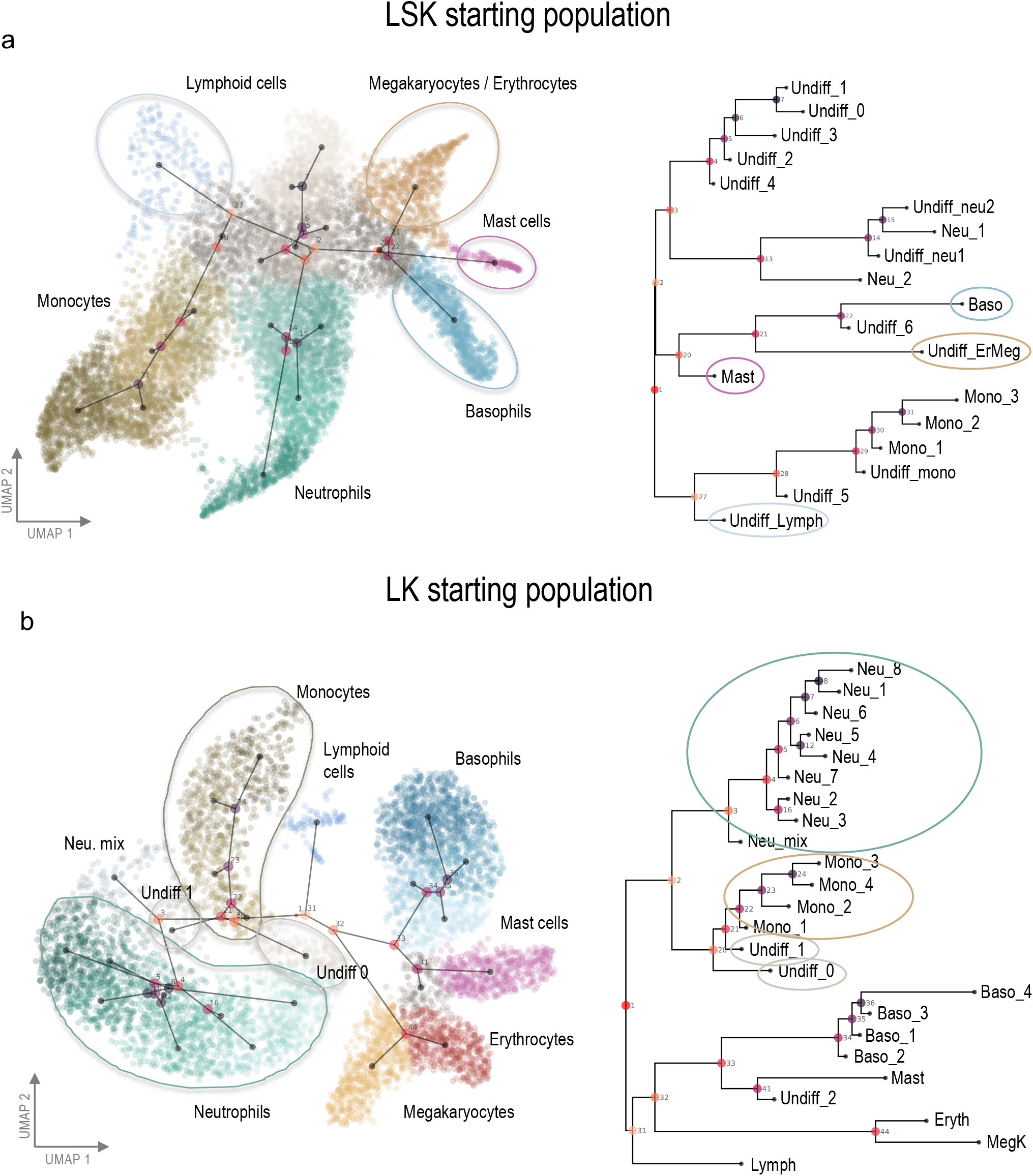
Application of PhyloTrajectory to lineage barcoding LARRY dataset. **a.** PhyloTrajectory analysis of hematopoietic cells from the LSK (Lin^−^Sca1^+^Kit^+^) starting population (well 2). UMAP colored by cluster identity with the inferred consensus trajectory projected onto the embedding (left) and the corresponding HIPSTR consensus tree (right). Circled groups on the tree highlight selected lineage branches. **b**. As in **a**, for the LK (Lin^−^Sca1^−^Kit^+^) starting population (well 2).

Several single-cell transcriptomic analyses have revised the classical organization of myeloid progenitor differentiation by placing basophil- and mast-cell-associated programs closer to the erythroid–megakaryocyte axis than to the neutrophil/monocyte branch [34], [35], [36]. Consistent with this, phylotrajectories reconstructed from the LK and LSK-derived population placed basophil and mast-cell clusters on the erythroid– megakaryocyte arm, and this is supported by the barcode correlation patterns, although the specific branching order (see e.g. the placement of mast cells) was not always consistent (Figure 5, a,b; Supplementary Figure S8 & S9).

The classical hematopoietic hierarchy posits an early bifurcation into common lymphoid progenitors (CLPs) and common myeloid progenitors (CMPs). Subsequent work, however, challenged this strict lymphoid–myeloid split by defining lymphoid-primed multipotent progenitors (LMPPs), which retain lymphoid together with granulocyte–monocyte potential after loss of erythroid–megakaryocyte potential [37], [38]. In the LSK-derived phylotrajectories, the lymphoid-associated state was positioned on the monocyte-associated side of the tree rather than branching from a node ancestral to both monocyte and neutrophil fates (Figure 5, a; Supplementary Figure S8).

In the LK-derived population, which is expected to be more committed upon barcoding, the lymphoid lineage branches off independently very close to the root (Figure 5, b), with a correspondingly low clade confidence (less than 0.7 for both well replicates), suggesting that the barcoding was after lymphoid divergence for this population.

Also for the LK starting population, the branching structure of the trees clearly groups monocytes and neutrophils, consistent with cells transiting through a granulocyte–monocyte progenitor (GMP)-like state [39], [33]. While no sampled cell subsets are closely associated with this progenitor, there are sampled states within this monocyte/neutrophil sub-lineage that appear to be transcriptionally undifferentiated. There are also transcriptionally undifferentiated states (e.g. Undiff_1) that, in dimensionality-reduced transcriptional space, are positioned distal to the central undifferentiated population (see Figure S7 B). Undiff_1 appears relatively basal to neutrophils (well 2, Figure 5, b) or monocytes (well 1, Figure S9, a), consistent with its undifferentiated state.

## Discussion

PhyloTrajectory introduces a phylogenetic framework for inferring T cell differentiation hierarchies using TCR sequences as endogenous barcodes. By modeling clonotype frequency evolution as a continuous stochastic process, the method recovers cellular state relationships from shared clonal patterns without requiring temporal sampling or experimental lineage tracing.

The core principle of PhyloTrajectory is that differentiation relationships between cellular states are encoded in clonotype frequency covariation. If two states are connected through a recent branching event, clonotypes that expanded before or during this transition will tend to appear in correlated abundances in both states; states separated by deeper or independent differentiation events will share weaker and more diffuse clonal structure. Clonotype frequency covariance thus acts as a statistical proxy for hierarchical relatedness.

A similar principle was exploited in a method by Weinreb and Klein (2020), showing that under a tree-structured branching-process model with suitable assumptions on differentiation dynamics, pairwise normalized covariances of clone counts can inform lineage topology and support hierarchy reconstruction from barcode data [40]. Their agglomerative merge-based algorithm reconstructs tree topology, but without any notion of branch lengths.

PhyloTrajectory relies on the same principle, but uses a full probabilistic model over the joint distribution of clonotype counts for each cell subset. This affords PhyloTrajectory a notion of branch length, which corresponds to accumulated clonotype frequency changes along a branch. In phylogenetics, branch lengths are critical for distinguishing deep structured clear divergences from shallow divergences that separate relatively similar states, and for representing and inferring multifurcations in binary trees (using internal branch lengths of zero). An explicit probabilistic model also allows us to sample from the posterior distribution over trees, here using MCMC. As with the three LCMV replicates, this can be used to adjudicate whether the model is asserting a statistically confident difference between processes, or if an observed difference could be due to sampling variation.

Any continuous process can be used in this approach, so long as likelihoods can be computed or approximated with enough stability to allow tree and parameter search and/or sampling. We used the Ornstein-Uhlenbeck process, which represents the simplest continuous process with an equilibrium distribution capable of capturing the essential dynamics of clonal evolution during differentiation: clonotype frequencies fluctuate stochastically while experiencing homeostatic pressures that prevent unlimited expansion. Just like nucleotide models used successfully in phylogenetics, this is intended as a mathematically tractable means of capturing the correlational structure in the observations induced by the branching tree topology, rather than a precise description of biological reality. We expect that, just as in traditional phylogenetics, tree topology inference will be relatively insensitive to the choice of continuous process.

Within this framework, the tree encodes the unobserved differentiation hierarchy, and branch lengths quantify the extent of clonal divergence between states. Notably, the model does not explicitly parameterize clonotype-specific lineage bias. Instead, lineage preferences are accommodated implicitly: preferential expansion along one branch (and possible contraction along another) manifests as stochastic frequency shifts under the OU dynamics. While explicit modeling of lineage bias could be introduced, statistical power to detect such effects at the level of individual clonotypes would likely be limited given typical sample sizes.

In single-cell clonotype analyses it might be tempting to observe a single clonotype that is particularly expanded in one cellular subset and, from this alone, conclude that there must have been some sort of functional reason, possibly driven by the TCR, for this apparent association. Outside of inferring differentiation topologies, our approach can serve as a conceptual null model that clarifies that, in the complete absence of any functional biases, such patterns would be expected from stochasticity alone, and any test for systematic biases needs to take this into account.

Application to LCMV infection data yielded differentiation hierarchies consistent with established models of CD4^+^ T cell responses. Notably, the IFN-stimulated cluster localized along the internal backbone upstream of terminal effector-associated states. Its transcriptional profile and central topological position support the interpretation of this population as a transient, plastic progenitor state [41]. In our inference framework, proximity to the root reflects broad, relatively unexpanded clonal structure, consistent with an early, multipotent phase preceding lineage commitment.

Effector populations (Th1 and Tfh) occupied distal branches, reflecting clonal specialization and expansion following lineage commitment. Although the global structure was reproducible across the three mice, the posterior consensus trees differed in the specific distal effector arrangement. Here, the model’s quantification of confidence was low in this region of the tree, indicating that this difference may just reflect inferential uncertainty due to sampling noise and limited data, rather than a genuine biological difference.

In both HDM and dog allergen models, naïve populations localized at the root and effector populations at terminal branches, indicating consistent global differentiation architecture. However, Treg positioning differed markedly between conditions and the frequency of Treg in the HDM model was substantially higher than that observed in the dog allergen model. In the initial analysis of the HDM model, Tregs formed a relatively long and distinct branch adjacent to Th2 populations. Clonal clustering revealed subsets of clones shared between Treg and Th2 cell states.

In contrast, in the dog allergen model, Tregs were positioned closer to the root with shorter branch lengths, indicating limited clonal divergence and minimal effector-associated expansion. This topology suggests a more central, less polarized regulatory phenotype. The dog allergen dataset also displayed polyclonal, cross-state clonal dynamics spanning Th17 and Th2 branches.

The long Treg branch in HDM prompted deeper investigation of internal heterogeneity. Re-clustering of the HDM Treg population revealed four transcriptionally distinct subsets that occupied non-overlapping positions along the inferred trajectory. A Helios^+^Cd27^+^ subset localized closest to the root and showed minimal clonal expansion, consistent with a stable regulatory population, possibly thymic-derived [27], [28]. In contrast, the remaining subsets—including an IFN-responsive, Gzmb-expressing population and a Foxp3-low Ccr5^+^ subset—were positioned further downstream and shared clonotypes with Th2 effector branches, indicating greater clonal expansion and effector-associated dynamics. Together, these findings indicate that the Treg compartment in HDM is not a single terminal state but a clonally stratified lineage spanning multiple differentiation niches [42]. Root-proximal Tregs appeared clonally unexpanded, whereas distal subsets displayed expansion patterns consistent with context-dependent functional adaptation within the Th2-polarized response.

Application to the LARRY lineage barcoding dataset [33] demonstrated that the method applies to datasets beyond endogenous T-cell receptor barcoding. Reconstructed trees largely recapitulated canonical hematopoietic organization, including coherent megakaryocyte–erythroid and granulocyte modules [43], [44].

The assumption of a strictly tree-structured topology reflects a hierarchical model of differentiation with progressive fate restriction. While appropriate for many systems, some states may arise via convergent or plastic pathways that violate strict tree structure. As shown, regulatory T cells, for example, may originate via both thymic and peripheral routes. If these are unable to be resolved via further subclustering, such scenarios may require network-based generalizations beyond tree topologies.

Several technical factors constrain inference. First, the signal driving inference is the number of expanded clones included in a sample, playing a similar role to the number of non-invariant nucleotide sites in traditional phylogenetics. Experimental factors that reduce sampling of expanded clones (low sampling depth, poor TCR recovery, etc) will reduce the quality of the inferred trees. Similarly, systems that just do not have many expanded clones will not be amenable to study by this approach.

Second, the temporal persistence of the signals required for correct topology inference is unclear. The method appears to perform well during active immune responses, but it may degrade in systems with long-running T-cell development spanning multiple years.

Third, the approach depends on gene expression–based clustering to define cellular states and, as shown with the Treg HDM example, overly coarse clustering can mislead inference. Clustering too finely, on the other hand, will reduce the number of clones per cluster and increase the noise in the resulting inferred tree topology.

PhyloTrajectory establishes that an explicit probabilistic model of clonotype frequency can be used to reconstruct differentiation tree structure from clone counts. Demonstrated with TCRs as natural endogenous lineage markers, as well as experimentally introduced LARRY barcodes, we anticipate that it will be extensible to other systems with differentiating cell subsets and lineage markers, including B-cells and other lineage barcoding systems. By reconstructing quantitative phylogenetic relationships from clonotype frequency patterns, the method offers a complementary perspective to expression-based trajectory inference, grounded in heritable clonal relationships rather than transcriptional similarity alone.

In traditional phylogenetics, sequence changes arise from the aggregate behavior of individual genotypes undergoing mutation, followed by frequency fluctuations that sometimes result in fixation of mutations. Phylogenetics approximates this with a species-level substitution process. Similarly, clonotype count distributions arise from a cell-level process. Here we have avoided an explicit description of the cell-level processes and instead directly specify the clonotype-level description of the distribution induced by a given tree. Future work will explore a range of cell-level processes, and their connection to the clonotype count distributions induced over bifurcating trees.

Future work will also aim to compare the behavior and performance of PhyloTrajectory compared to topology-only tree reconstruction methods [40] and expression-based trajectory inference approaches. Given that, to varying degrees, the assumptions behind these methods and the objects that are inferred by them differ, this will require a careful simulation study that is not unduly biased in favour of any specific method.

## Materials and Methods

### Clonotype count matrices

Phylotrajectory takes as input a clonotype count matrix *C* ∈ ℕ^*K*×*M*^, where *K* denotes the number of transcriptionally defined cell clusters and *M* the number of clonotypes. Cell clusters are inferred from gene expression data, while clonotypes are defined from TCR sequences or lineage-barcoding data using standard repertoire reconstruction pipelines or by direct sequence identity. Only cells with assigned clonotypes are retained. Each matrix entry *C*_*k,m*_ corresponds to the number of cells of clonotype *m* observed in cluster *k*.

### PhyloTrajectory Model

We describe the relationship between cellular states with a differentiation tree. Each leaf node in the tree is associated with an observed cellular state, and the internal nodes and branches on the tree represent transient differentiation events and transitions between states.

Our model considers each clonotype independently. At the root of the tree a clonotype is typically at low frequency, and propagates from the root to the tips changing in frequency along the branches. At a branching event, the frequency of the clonotype is inherited along the child branches. Thus the differentiation tree induces correlational structure in the joint distribution of the clonotype frequencies of the leaf nodes.

Specifically, for a given clonotype *j*, we model the log-frequency *F* using an Ornstein–Uhlenbeck (OU) process. This combines Brownian diffusion with a restoring force towards an equilibrium mean frequency *μ*:

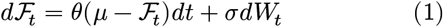

where *W*_*t*_ is a Wiener process, *σ* (volatility) controls the magnitude of log-frequency fluctuations (expansion and decay), and *θ* (*θ* > 0) is the mean-reversion rate and controls how strongly the process is pulled back towards the equilibrium mean *μ*, representing a homeostatic effect that prevents runaway expansion of a single clonotype.

As in phylogenetics, there is a confound between the process rates and total tree length (ie. the sum of all branch lengths), so we fix *σ* = 1 which sets the unit of branch length to ‘diffusion time’: for short branches, variance accumulates at approximately one unit per unit branch length (and exactly so under the Brownian limit *θ* → 0). For familiarity we will include *σ* in our description.

We model the initial log-frequency at the root of the tree with a Gaussian, with mean *μ*_*R*_ and standard deviation

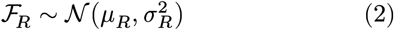

Note: to match the biology where the early pre-differentiated clonotype log-frequencies are typically low with low variance, our root state is not tied to the equilibrium distribution of the process under which the log-frequencies evolve. As such, this process is non-stationary and the likelihood is not invariant to the position of the root on the tree (as is the case in many standard models of molecular evolution).

At each leaf node *i*, we observe clonotype counts *C*_*ij*_ for clonotype *j*. These are modeled as Poisson-distributed realizations whose rate parameter depends on the unobserved, latent log-frequency *F*_*ij*_:

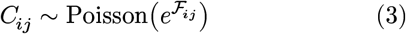

The generative model that induces a distribution over the clonotype counts can thus be described by four parameters, *θ, μ, μ*_*R*_ and *σ*_*R*_, the tree topology, and the branch lengths.

### Prior distributions in the model

We adopt a Bayesian framework, placing prior distributions on all model parameters to complete the probabilistic specification. The priors are chosen to be weakly informative, allowing the data to dominate inference while ensuring proper posterior distributions, with the exception of the prior on the root state (see below) which is intended to be informative.

#### Branch lengths

Branch lengths are constrained to be positive and are parameterized on the logarithmic scale. We place a Gaussian prior on the log branch length *l, l* ∼ *N* (−1.0, 1.0). This form ensures positivity while allowing for occasional longer branches.

#### Root state

The log-frequency at the root node for each clonotype is modeled with a Gaussian prior 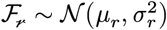, where both the mean *μ*_*r*_ and 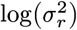 are themselves assigned Gaussian priors *N* (0, 1.0) and *N* (0, 0.1) respectively, reflecting the expectation that ancestral clonotypes begin at low frequencies with limited initial variance.

#### Root position

The position of the root along the tree is sampled using a uniform prior over the total branch length of the whole tree.

#### Equilibrium mean

The equilibrium mean *μ* toward which log-frequencies revert is assigned a Gaussian prior: *μ* ∼ *N* (1.5, 1.0), reflecting a prior expectation of moderate clonotype frequencies at equilibrium.

#### Mean-reversion rate

The mean-reversion parameter *θ* is sampled on the log scale to ensure positivity. A Gaussian prior is placed on log(*θ*): log(*θ*) ∼ *N* (−1.0, 1.0) allowing the data to determine whether clonotype dynamics are dominated by drift (small *θ*) or mean-reversion (large *θ*).

### Likelihood calculation

Under the OU process governing log frequencies, the transition density between a parent node *p* and child node *i* with branch length *t*_*i*_ is Gaussian:

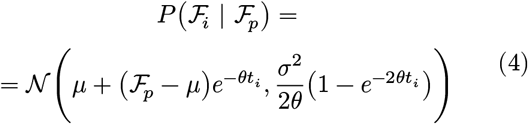

Leaf counts *C*_*ij*_ are Poisson 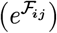, but efficient likelihood calculation for continuous states requires Gaussian sub-tree likelihoods. For large counts, the Poisson likelihood *P* (*C*_*ij*_ | *F*_*ij*_) (i.e. considered as a function of *F*_*ij*_) is well approximated by a (scaled) Gaussian density with moments:

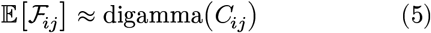

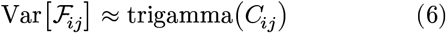

Inference only requires the un-normalized likelihood so the scaling constants for this approximation are not required. For lower counts, this approximation is weaker, and when *C*_*ij*_ = 0 the likelihood *P* (*C*_*ij*_ | *F*_*ij*_) is monot-onically decreasing and has no left tail, and so we use a pseudocount of 0.5 when approximating *P* (*C*_*ij*_ | *F*_*ij*_) for *C*_*ij*_ = 0. The robustness of inference under this approximation is justified by the simulation study where no approximation was used generating the data, but inference used this Gaussian leaf likelihood approximation.

Given a tree *T* with internal node indices 1, ‥, *n* (where the root is 1) and leaf node indices *n* + 1, …, *l*, the latent log-frequency at node *i* for clonotype *j* ∈ {1, …, *m*} is *F*_*ij*_ ∈ ℝ, and *C*_*ij*_ ∈ ℕ_0_ the observed count at leaf *i*.

The likelihood is then

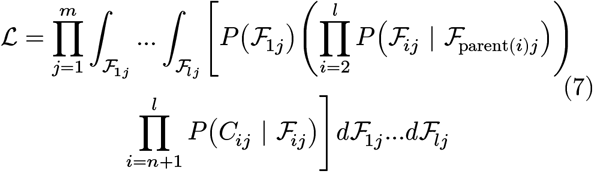

Naively, this integral would be intractable. However, when the likelihood of the counts for a single subtree, marginalizing over the subtree internal nodes, can be represented as a re-scaled Gaussian density, then it can be rearranged using exactly the same strategy as in Felsenstein’s pruning algorithm [14] and calculated in time linear in the number of counts. With the Gaussian leaf likelihood approximation and Gaussian transition density under the OU process, it suffices to note that the pointwise product of two scaled Gaussians is another Gaussian, and thus all of the sub-tree likelihoods are Gaussian. This is implemented in `MolecularEvolution.jl` [45] (see below).

### Model inference and convergence

Posterior inference over the model parameters and tree configuration is performed using a composite MCMC sampler combining Gibbs and Metropolis–Hastings updates.

At each sampling step, the following parameters are sequentially updated:

- the tree topology, via a Gibbs sweep over all nearest-neighbor interchanges, selecting a topology *T* ^′^ proportional to its posterior probability,
- branch lengths, via a Gaussian random-walk on the log branch length with proposal distribution *N* (0, 0.1^2^), yielding a multiplicative proposal on the branch length scale,
- the root state, representing the ancestral clonal abundance at the origin of the tree, is updated using a Gaussian random-walk proposal with mean and log variance represented by *N* (0, 0.1^2^) and *N* (0, 0.1^2^). The root position is jointly updated with state via proposals drawn from a local uniform distribution within a radius proportional to the total tree length (default: 10% of total branch length).
- Proposals on the equilibrium mean *μ*, defining the long-term average clonotype frequency, the mean-reversion rate *θ*, determining the rate of stabilization towards *μ*, are drawn from *N* (0, 2.0^2^) and *N* (0, 0.5^2^) respectively.

The topology update selects among competing interchanges in a single Gibbs step, while the remaining continuous parameters are accepted or rejected according to the standard Metropolis–Hastings rule. Topology and branch length proposals are ordered for efficient evaluation via traversals of the tree that locally re-propagate messages after any change, ensuring likelihood consistency with only constant-time updates per evaluated proposal.

The tree is initialized as a random bifurcating phylogeny with number of leaves equal to the number of gene expression clusters, effective population size *N*_*e*_ = 1.0, sample rate equal to 100 and starting branch length 0.1. Before the main sampling phase, an optional warmup performs iterations (default: 100) updating only the tree topology and root state and position, with branch lengths and process parameters held fixed, allowing the tree and root configuration to reach a reasonable starting point before joint inference begins.

Representative post-burn-in MCMC trace plots for the LCMV and LARRY LK-subset analyses, including LARRY LK wells 1 and 2, are provided in Supplementary Figure S10. These plots were used as posterior-sampling diagnostics to assess sampling stability after burn-in and to check that posterior summaries were not dominated by transient early-chain behavior.

### Implementation

`MolecularEvolution.jl` offers a set of generically implemented algorithms that require, for a new process, implementation of the local behavior along a branch, including the “forward time” transition density from parent to child and reverse time likelihood propagation from child to parent, as well as the combination of evidence from two conditionally independent subtrees [45]. This affords Felsenstein’s upward (leaf to root) message passing algorithm for likelihood calculations, a downward message-passing algorithm that aggregates evidence from all observations on the tree to every node, allowing for marginal state calculations and enabling efficient (constant time) likelihood re-evaluation under local tree rearrangement, such as branch length modification and nearest-neighbour interchange. It also supplies routines for branch length and tree topology optimization. For this paper, in `MolecularEvolution.jl` the OU process was implemented and we expanded the inference support to include Bayesian inference by Markov Chain Monte Carlo, including proposals to the root state and position to allow for efficient inference under non-stationary models.

### Simulations setup

Synthetic datasets were generated by propagating clonotype frequencies along a specified tree under a branch-wise stochastic model. Each simulation started from a random bifurcating tree with *N* leaves (4– 10). The sampling parameters were chosen to sample diverse tree structures (Supplementary Figure S11). After generation, total branch length was rescaled to 10.0 units to control the expected degree of clonal expansion across replicates.

Frequencies at the root were initialized from a Gaussian distribution, and branch-wise evolution followed a Brownian-motion process:

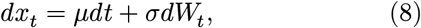

with mean drift *μ* = −0.3 and variance *σ*^2^ = 1.5, chosen to approximately recapitulate clone size distributions. Counts were sampled from a multinomial distribution, drawing *N* cells with probabilities given by the softmax of the log-frequencies at the leaves of the tree. Two settings were considered: a small setting (mean *N* of 15,000 cells per tree) and a large setting (mean *N* of 1,050,000 cells per tree). The number of clonotypes was fixed at 1500; because counts arise from multinomial sampling, however, some clonotypes may receive zero counts and be absent from a given matrix, so the realized number of observed clonotypes per tree is at most 1500.

For inference, each simulated count matrix was analyzed with the OU model described above. Each run used a warm-up phase of 100 trees, followed by 10,000 burn-in iterations and 1,000 posterior samples with a sampling interval of 10. At each step, the MCMC updated jointly proposed changes to tree topology, branch lengths, and OU parameters via a composite sampler described above.

For each replicate, the reconstructed tree was compared with the ground-truth tree using two metrics: (i) the Pearson correlation of tree distances from the true and inferred tree, where these distances include all pairwise distances between any two leaves, as well as the distance from the root to each leaf (the latter included so that the metric is sensitive to the accuracy of the inferred root position), and (ii) the number of unmatched internal nodes, where we consider nodes to match between two trees if they induce the same split upon the leaves. Each experiment was repeated 100 times for each number of leaves to assess robustness as tree complexity increased.

### Clone clustering

To identify patterns of evolutionary diversification of clonotypes across differentiation states, we performed clonal clustering based on OU-inferred marginal clone activities. Posterior samples of trees and Ornstein–Uhlenbeck (OU) parameters were obtained via a Metropolis–Hastings MCMC procedure as described above. From these samples, a representative consensus topology was derived using HIPSTR, and the median OU parameters across the chain were used to compute node-wise marginal posterior means of the latent clone log-frequencies. Each marginal distribution, defined by mean (*μ*) and variance (*σ*^2^), was converted to the expected abundance in count space using the transformation exp 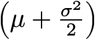, and these expected abundances were used for downstream clustering.

To detect clonally related groups with similar dynamic trajectories, we performed z-score normalization of each clone expected abundance vectors across all tree nodes and computed pairwise similarities. Clones were then clustered using the Dirichlet-process-means [21] algorithm—an adaptive, non-parametric extension of k-means that automatically determines the number of clusters based on a specified distance threshold (implemented in DPMeansClustering.jl, https://github.com/MurrellGroup/DPMeansClustering.jl [22]). This approach grouped clones exhibiting similar profiles across the differentiation landscape.

### Statistical test for sub-structure within clusters

To determine whether individual transcriptional clusters exhibited clonal size heterogeneity that was associated with their expression state, with an association that is stronger than expected by chance, we performed a nearest-neighbor–based permutation test on the PCA gene expression space. We are specifically interested in the expression-space distribution of the long right tail of the clonotype size distribution, and we want to ask: are there regions of PCA space that systematically have more of the larger clones than other regions? We constructed a test statistic that associates each cell with the maximum clonotype count of clonotypes associated with that cell’s k nearest neighbours (k=50) in PCA space. Our test statistic is the variance of this “local neighbour maximum” over the entire cluster.

To evaluate whether the observed variance exceeded what would be expected under the null hypothesis that cells (and their clonotype sizes) are uniformly distributed within a cluster, we performed 1,000 permutations in which clone labels were randomly shuffled within the cluster while preserving all cell coordinates in PCA space. For each shuffled cluster, the above statistic was computed to yield a null distribution. We then compared the observed test statistic with the empirical null distribution and calculated one-sided empirical permutation p-values, corresponding to the fraction of permuted statistics greater than or equal to the observed statistic. To account for testing multiple transcriptional clusters, p-values were adjusted using the Bonferroni-Holm procedure. An observed variance substantially exceeding the permutation distribution indicates that the cluster shows greater clonal peak size heterogeneity across PCA-space than expected by chance—that is, the cluster is not uniform, and contains unusually strong local regions with higher frequencies of expanded clones.

### Experimental animals

Wild-type C57BL/6J mice were bred and maintained at the Comparative Medicine animal facility at Karolinska Institutet. Mice were 6–8 weeks old at the start of experiments. Animals were housed in individually ventilated cages under specific pathogen-free conditions, with ad libitum access to food and water. All procedures were approved by Stockholms Jordbruksverket (permit numbers 8971/2017).

### House Dust Mite Model

Six-to eight-week-old mice were briefly anesthetized with isoflurane and sensitized intranasally with 1 μg house dust mite (HDM) extract in 40 μL PBS. Seven days later, mice received intranasal challenges of 10 μg HDM once daily for five consecutive days. Four days after the final challenge, mice were euthanized and tissues collected for analysis. Bronchoalveolar lavage (BAL) was performed by two sequential airway flushes with 1 mL PBS [46]. Lung tissue and mediastinal lymph nodes (medLN) were simultaneously harvested, but only the BAL compartment, representing the active T cell response, was used for this study.

### Single Cell RNA Sequencing

Samples were stained for surface markers and labeled with individual TotalSeq Hashtags (BioLegend) for sample identification of BAL (vs medLN and lung, simultaneously collected for another study). All samples were processed on ice to preserve cell integrity. Cells were stained with antibodies shown in Supplementary Table S3. T cells (CD4+ CD3+ B220−) and B cells (CD19+ B220+ CD3-, CD4-, also sampled for another study) were sorted into pure FCS using a BD FACSAria Fusion. Sorted cells were washed and resuspended in cold PBS, then processed for single-cell capture using the Chromium droplet-based microfluidic system (10x Genomics). Library preparation was performed by the Eukaryotic Single Cell Genomics national facility at SciLifeLab, Stockholm.

### scRNA-seq and scTCR-seq data processing

Data from Dog Allergen dataset (GSE244722) and house dust mite datasets were processed separately with cellranger-8.0.1 *multi* pipeline. Single-cell data analysis was performed using Scanpy (v1.11.1) under Python 3.11.8. During quality control (QC), cells with mitochondrial gene content exceeding 10% or fewer than 1000 total counts were excluded. Doublet detection was performed with Scrublet method implemented in scanpy; Cells with a doublet score below 0.2 were retained for downstream analysis. In total, 5,829 cells were left in dog allergen dataset and 6,331 cells in house dust mite dataset. The counts were further log-normalized. 3000 highly variable genes (HVGs) were selected, excluding those annotated as T cell receptor genes. Leiden clustering was performed on neighbors graph with parameters n_pcs=15 and n_neighbors=100. TCR clonotypes were obtained from cellranger filtered_contigs_annotation.tsv annotated file. The IFN-response cluster from DA model was excluded due to low number of cells with TCRs present. Proliferating clusters were excluded for inferring differentiation trees due to mixture of diverse populations.

### Cell state annotation in the LCMV dataset

Data from five mice from GSE158896 was used. QC of each dataset was performed using the same cutoffs as described above. Batch effects between mice were corrected using harmonypy with default parameter settings. Cell states were annotated by mapping clusters onto the LCMV CD4 T cell reference atlas using the ProjecTILs package in R [20]. TCR clonotypes were obtained from the cellranger filtered_contigs_annotation.tsv annotated file.

### LARRY lineage barcoding dataset

Lineage tracing data were obtained from GSE140802 [33]. We retrieved the normalized data from the *in vitro* experiment and reprocessed them to annotate clusters (Scanpy v1.11.1, Python 3.11.8). For each starting population (LSK and LK), we computed PCA on 3,000 highly variable genes and integrated across wells with the Harmony algorithm. The integrated principal components were used to build neighbor graphs and for subsequent clustering (n_pcs=20, n_neighbors=100). Leiden clustering at resolutions 1.5 (LSK) and 2.0 (LK) resulted in 24 and 26 clusters respectively. Cluster assignments were merged with barcode information, and cells lacking barcode data were removed. We then stratified the data by well (1 or 2) and removed clusters with fewer than 50 cells in either well due to insufficient signal.

## Data and code availability

The newly generated house dust mite (HDM) single-cell gene-expression and TCR-sequencing data generated in this study will be deposited and will be made publicly available upon publication. The previously published dog allergen, LCMV CD4^+^ T cell, and LARRY lineage-barcoding datasets used in this study are available under accessions GSE244722, GSE158896, and GSE140802, respectively. The PhyloTrajectory implementation and analysis scripts used in this study are available at https://github.com/MurrellGroup/Phylotr_ajectories.jl.

## Author contributions

M.G.: Conceptualization, Methodology, Software, Investigation, Formal analysis, Visualization, Writing – original draft. M.D.: Methodology and Software. C.A.T.: Investigation and Resources. J.C.: Supervision and Writing – review & editing. B.M.: Conceptualization, Methodology, Software, Supervision, Project administration, Funding acquisition and Writing – review & editing. All authors reviewed and approved the final manuscript.

---

**Figure S1.**
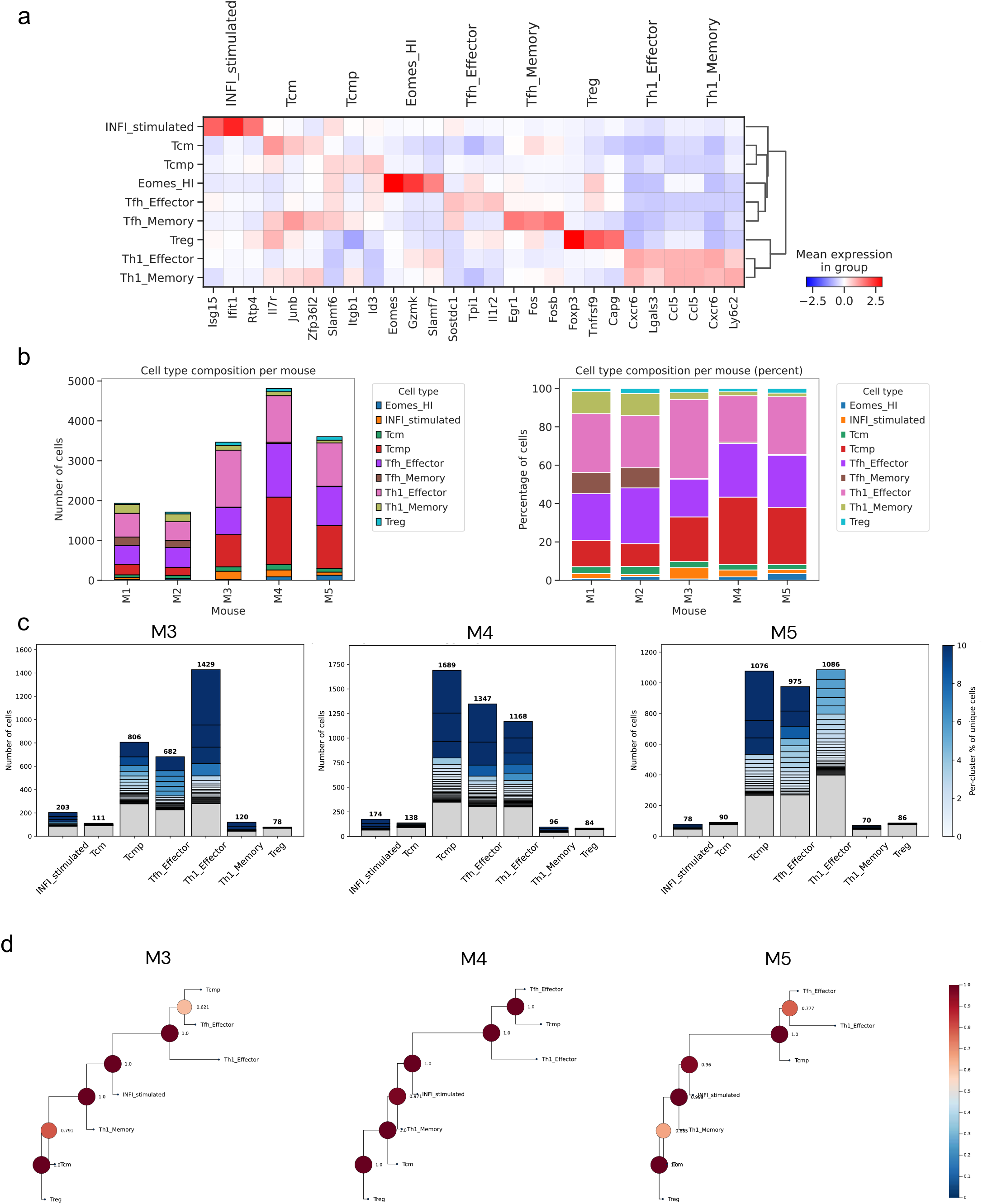
Cell type annotation and clonal composition in the LCMV dataset. a. Heatmap of marker gene expression across the annotated CD4^+^ T cell states identified in the LCMV Armstrong dataset, supporting cluster annotation via ProjecTILs reference mapping. Color scale represents mean expression within each group (z-scored). b. Bar plots show absolute cell counts (left) and percentage composition (right) per annotated cell state across the five replicates (M1–M5). c. Bar plots of absolute cell counts for the top 10% most expanded clonotypes per cell state for M3, M4, and M5 (left to right). d. PhyloTrajectory-inferred differentiation trees for M3, M4, and M5 colored by confidence scores.

**Figure S2.**
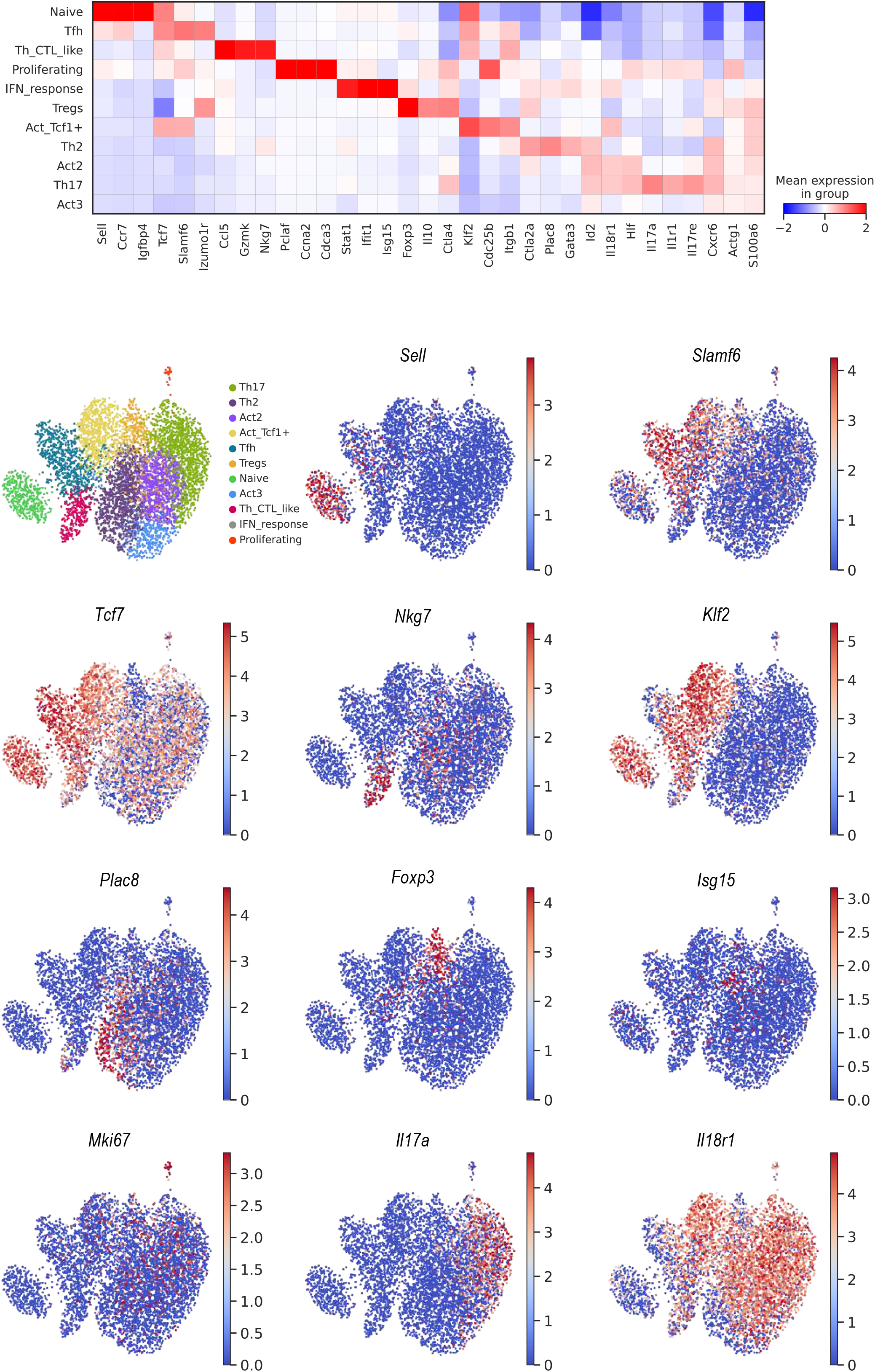
Cluster characterisation in the Dog Allergen dataset. Heatmap of top marker genes expression across the CD4^+^ T cell clusters identified in the Dog Allergen model. Color scale represents mean expression within each group (z-scaled). UMAP plots of Dog Allergen cells colored by cluster identity and by expression of key marker genes: *Sell, Slamf6, Nkg7, Klf2, Plac8, Foxp3, Isg15, Mki67, Il17a*, and *Il18r1*.

**Figure S3.**
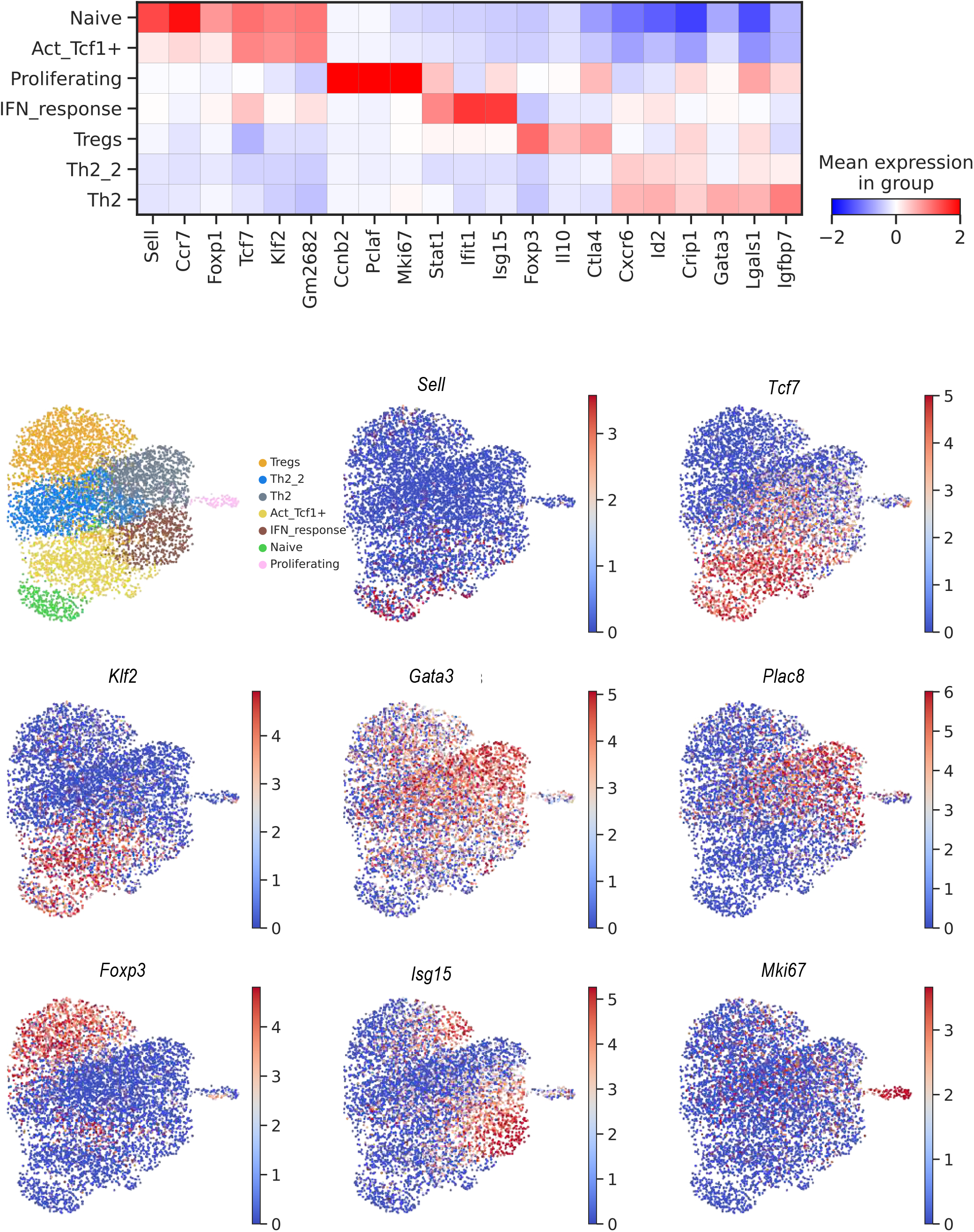
Cluster characterisation in the HDM dataset. Heatmap of top marker genes expression across the CD4^+^ T cell clusters identified in the HDM model, including activated circulating. Color scale represents mean expression within each group (z-scaled). UMAP plots of HDM cells, colored by cluster identity and by the expression of key marker genes used for annotation: *Sell, Tcf7, Klf2, Gata3, Plac8, Foxp3, Isg15*, and *Mki67* (bottom).

**Figure S4.**
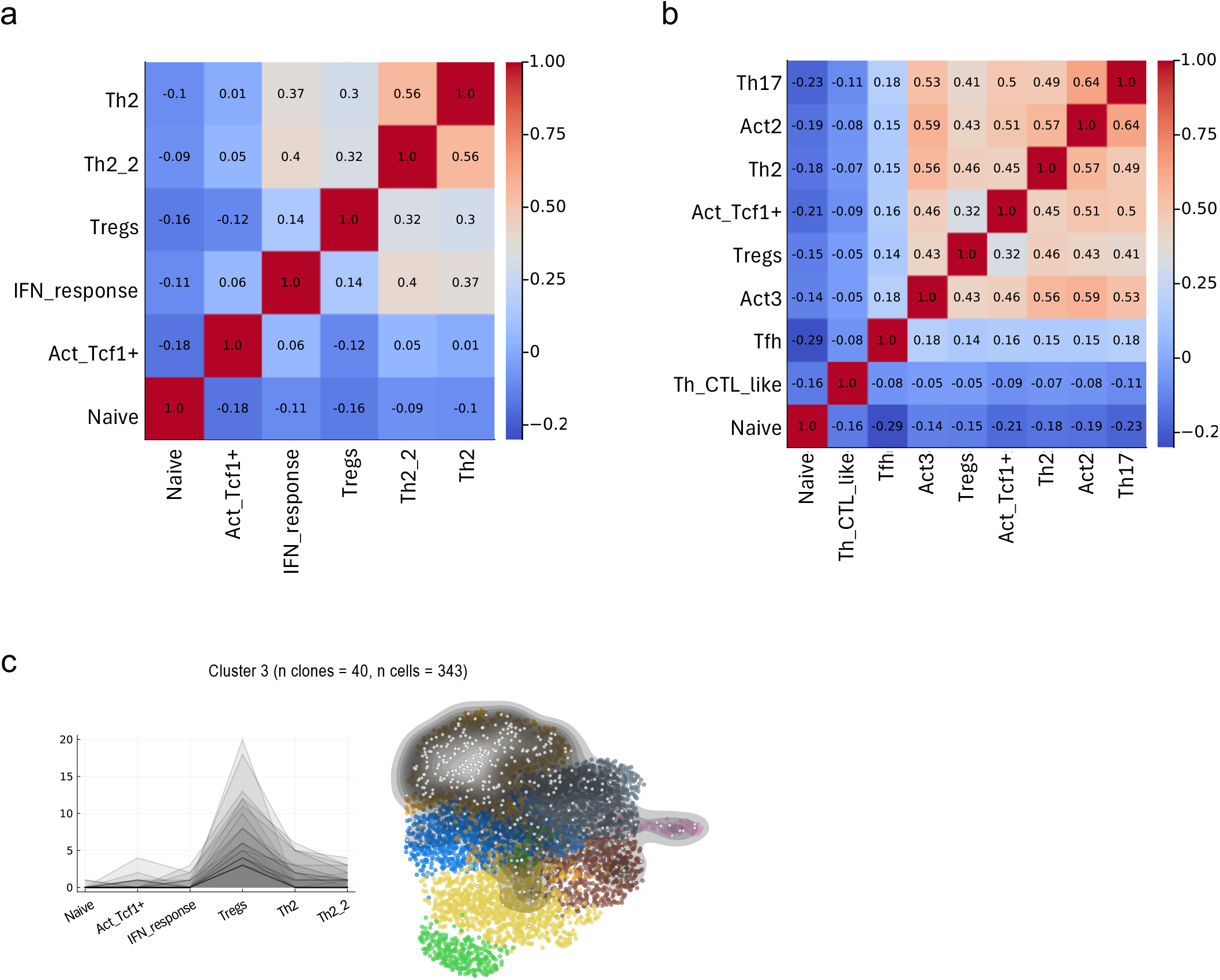
Clonotype frequency correlation heatmaps and representative clonal cluster in the allergen datasets. a. Pairwise Pearson correlation matrix of log-transformed clonotype counts across cell types in the house dust mite (HDM) model. b. As in a, for the Dog Allergen (DA) model. c. HDM clonal Cluster 3 (n clones = 40, n cells = 343), enriched in the Treg population. Individual clonotype cell counts across cell types are shown on the left and the UMAP density overlay on the right.

**Figure S5.**
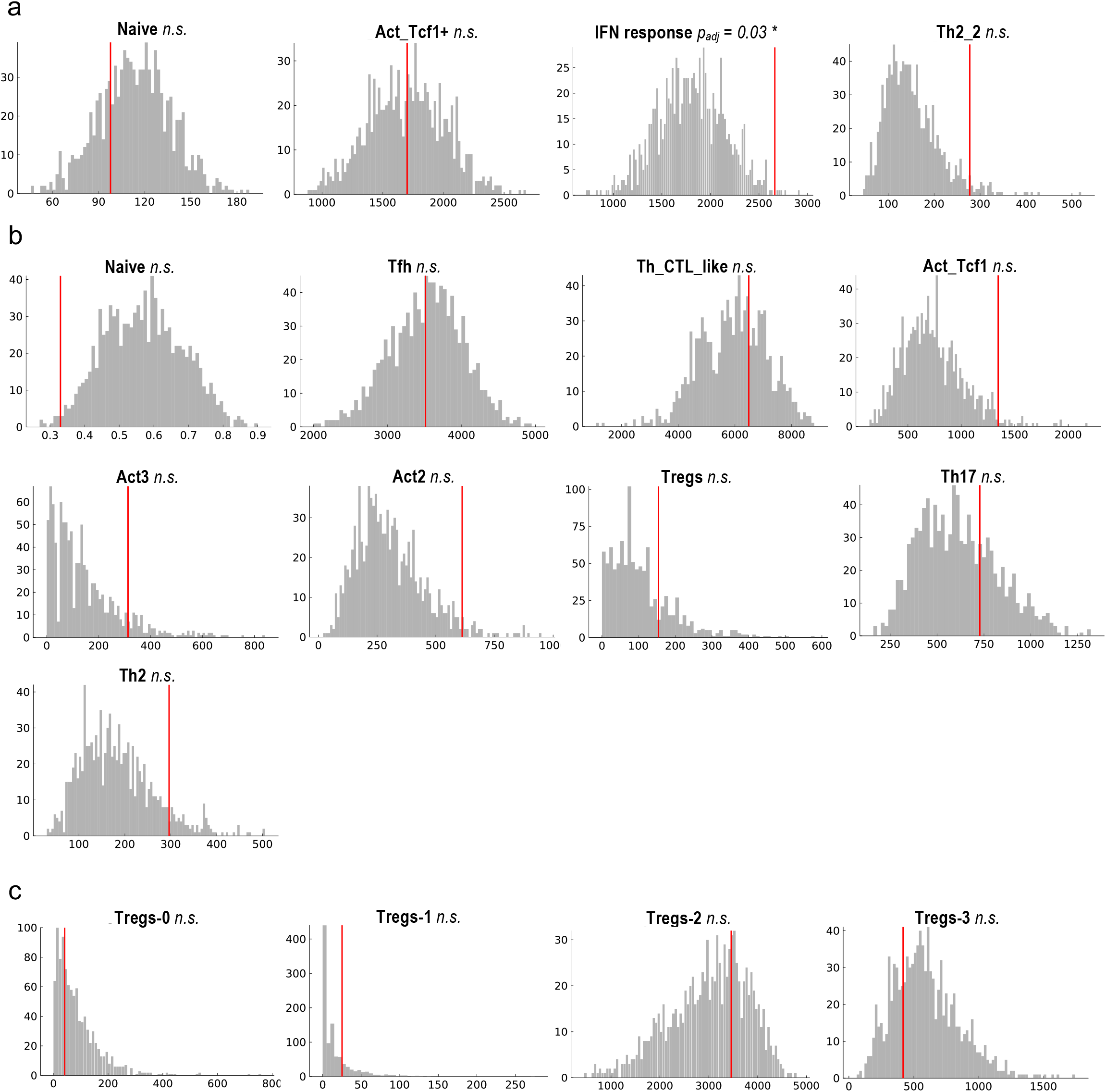
Additional permutation tests for local clonal size heterogeneity. a. Additional nearest-neighbour permutation test results for HDM transcriptional clusters not shown in Main Fig. 4a. For each cluster, the observed variance of the local-neighbourhood maximum clonotype size in PCA space is shown by the red vertical line and compared with the null distribution generated from 1,000 within-cluster clone-label permutations, shown as grey histograms. Empirical one-sided permutation p-values were adjusted across clusters using the Bonferroni-Holm procedure. b. Nearest-neighbour permutation test results for transcriptional clusters in the Dog Allergen dataset. The observed statistic is shown by the red vertical line and the permutation null distribution by the grey histogram. None of the tested Dog Allergen clusters showed significantly elevated local clonal size heterogeneity. c. The same permutation test applied to the four HDM Treg subpopulations, Tregs-0, Tregs-1, Tregs-2, and Tregs-3, following re-clustering of the original Treg population. None of the Treg subpopulations showed significantly elevated local clonal size variance relative to the permutation null, consistent with reduced residual clonal size substructure after Treg re-clustering. p_adj_, Bonferroni—Holm adjusted permutation p-value; n.s., not significant; *p_adj_ < 0.05; **p_adj_ < 0.01;

**Figure S6.**
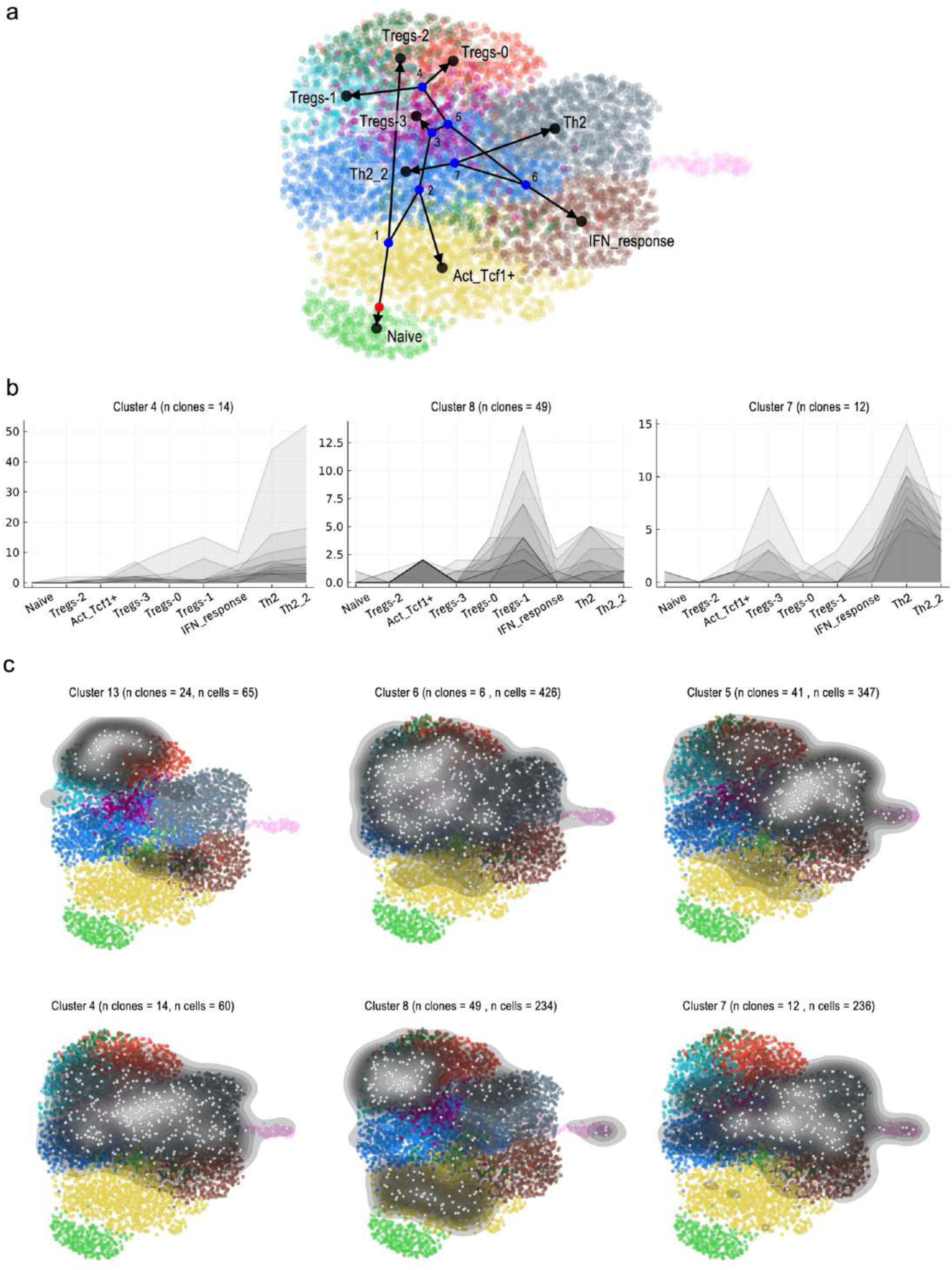
Extended clonal cluster characterization in the HDM dataset following Treg re-clustering. a. PhyloTrajectory-inferred differentiation trajectory overlaid on the UMAP embedding of HDM CD4+ T cells, with Treg subpopulations resolved into Tregs-0, Tregs-1, Tregs-2, Tregs-3. b. Line plots showing observed cell counts across cell states for three additional clonal clusters (Clusters 4, 8, and 7) not shown in the main figure. The number of clonotypes per cluster is indicated above each panel. c. UMAP density overlays showing the spatial distribution of clonotypes from six selected clonal clusters (Clusters 13, 6, 5, 4, 8, and 7). The number of clonotypes and total cells per cluster is indicated above each panel.

**Figure S7.**
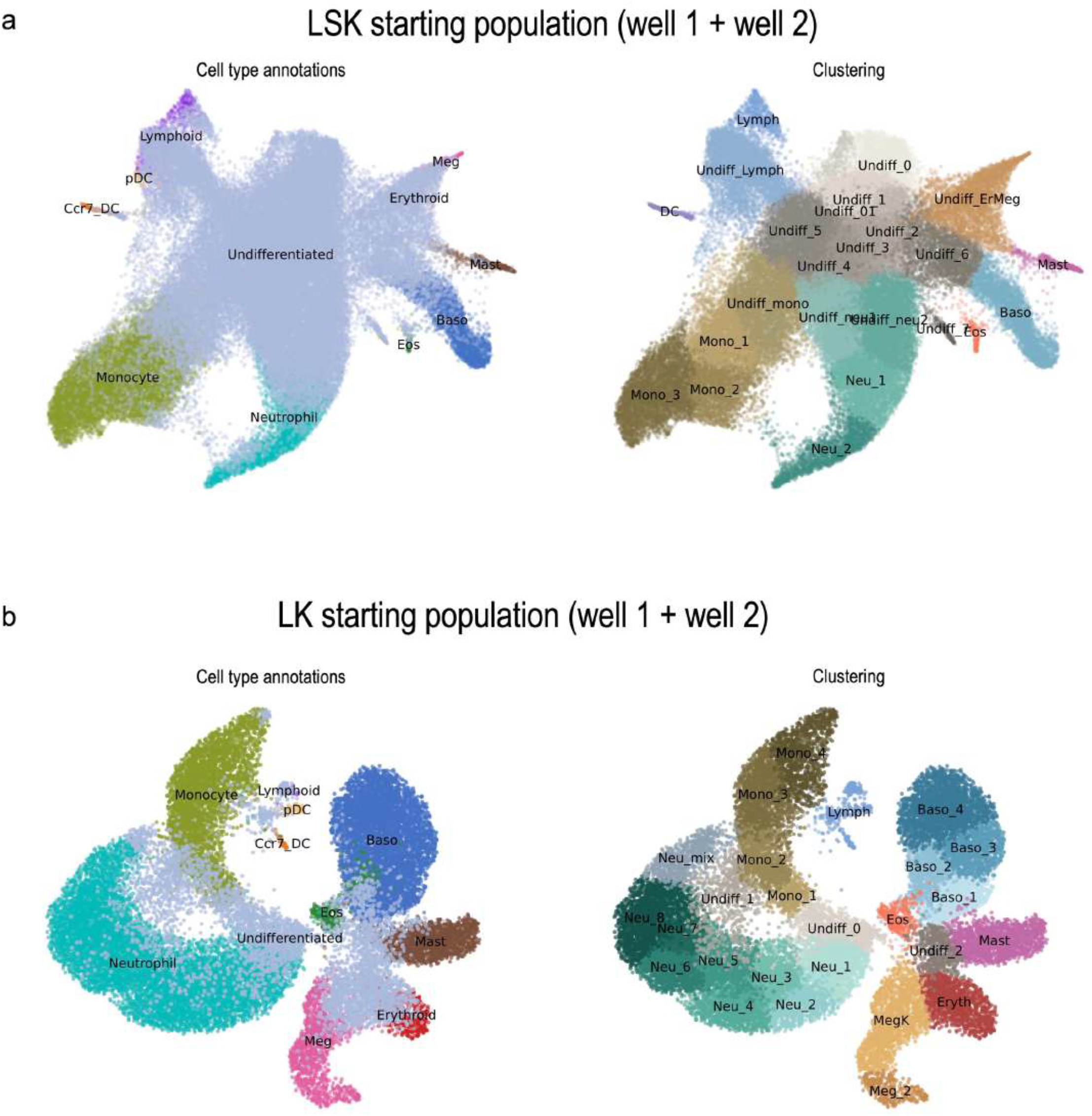
Cell type and clustering annotations for LARRY hematopoietic cells across starting populations. UMAP embeddings of cells from the two starting populations, with wells 1 and 2 pooled in each case. a. LSK (Lin^−^Sca1^+^Kit^+^) starting population. On the left, cells colored by the cell-type annotations from the original study Weinreb et al. (2020) [33]. On the right, the same embedding colored by the clusters used for the PhyloTrajectory analysis. b. LK (Lin^−^Sca1^−^Kit^+^) starting population, shown analogously to a: left, original cell-type annotations; right, clusters used for the PhyloTrajectory analysis

**Figure S8.**
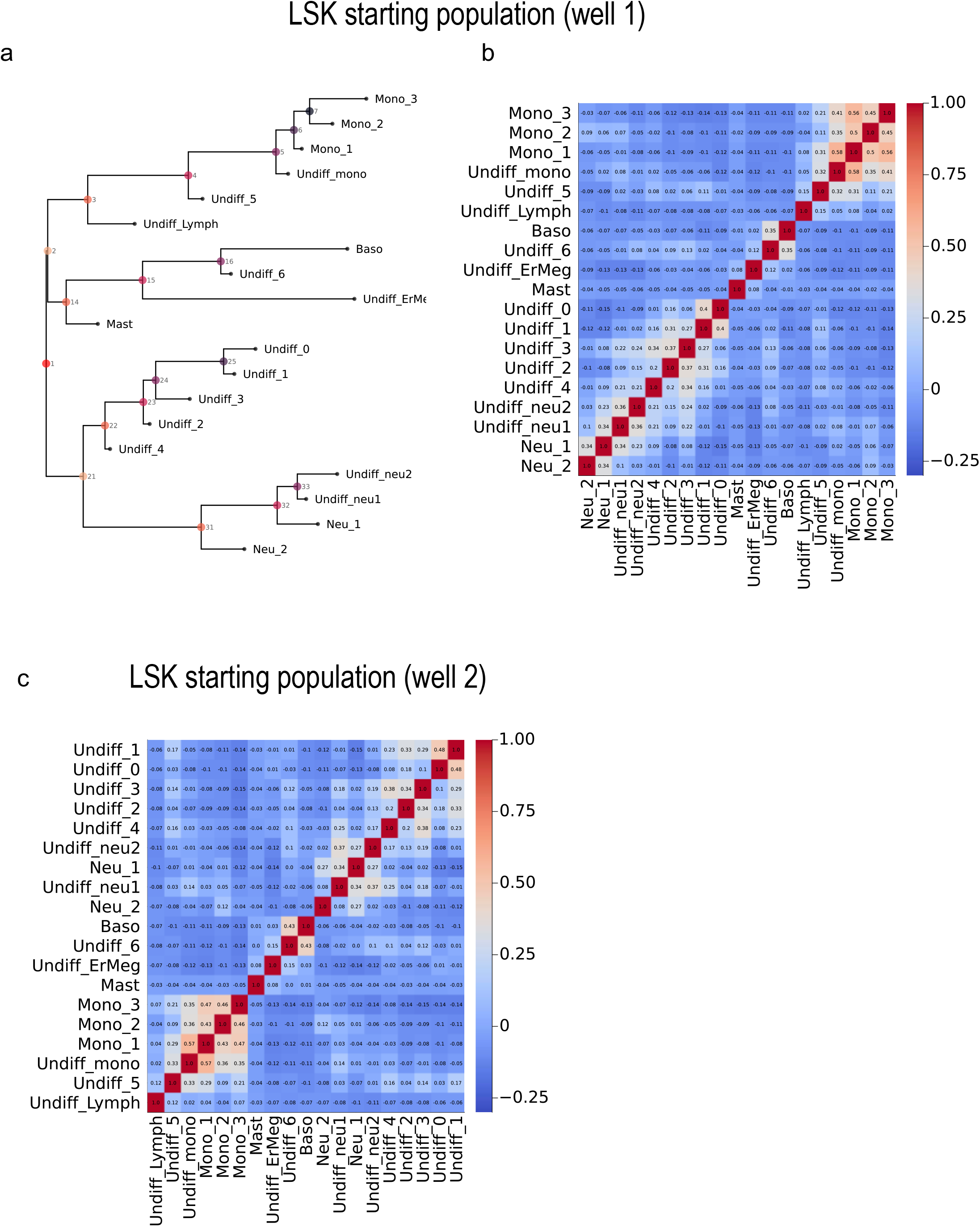
Barcode correlation heatmaps and PhyloTrajectory-inferred tree for the LSK starting population. a. PhyloTrajectory-inferred consensus tree for LSK well 1. b. Pairwise Pearson correlation matrix of log-transformed barcode counts between clusters for LSK well 1. c. Pairwise Pearson correlation matrix of log-transformed barcode counts between clusters for LSK well 2.

**Figure S9.**
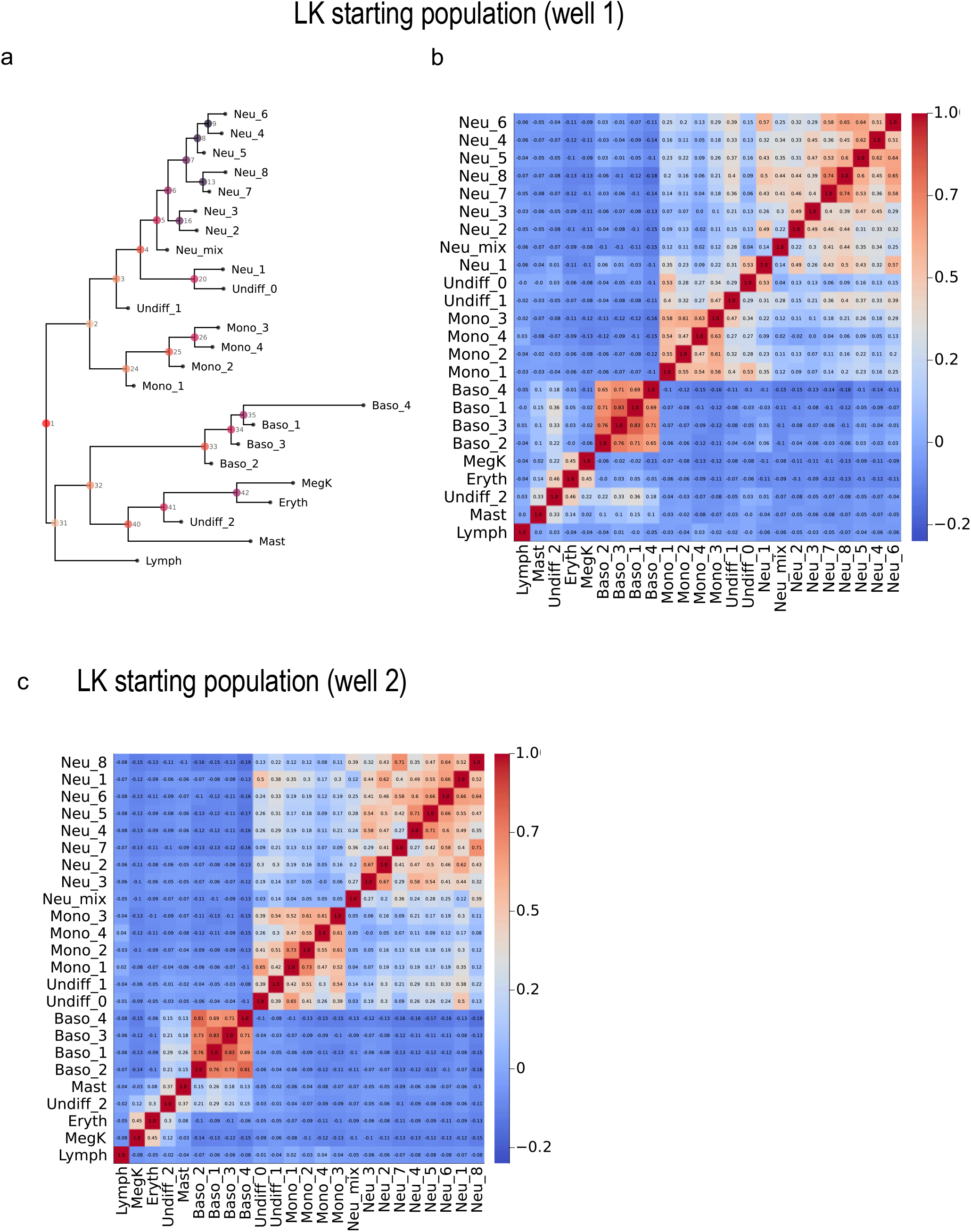
Barcode correlation heatmaps and PhyloTrajectory-inferred tree for the LK starting population. a. PhyloTrajectory-inferred consensus tree for LK well 1. b. Pairwise Pearson correlation matrix of log-transformed barcode counts between clusters for LK well 1. c. Pairwise Pearson correlation matrix of log-transformed barcode counts between clusters for LK well 2.

**Figure S10.**
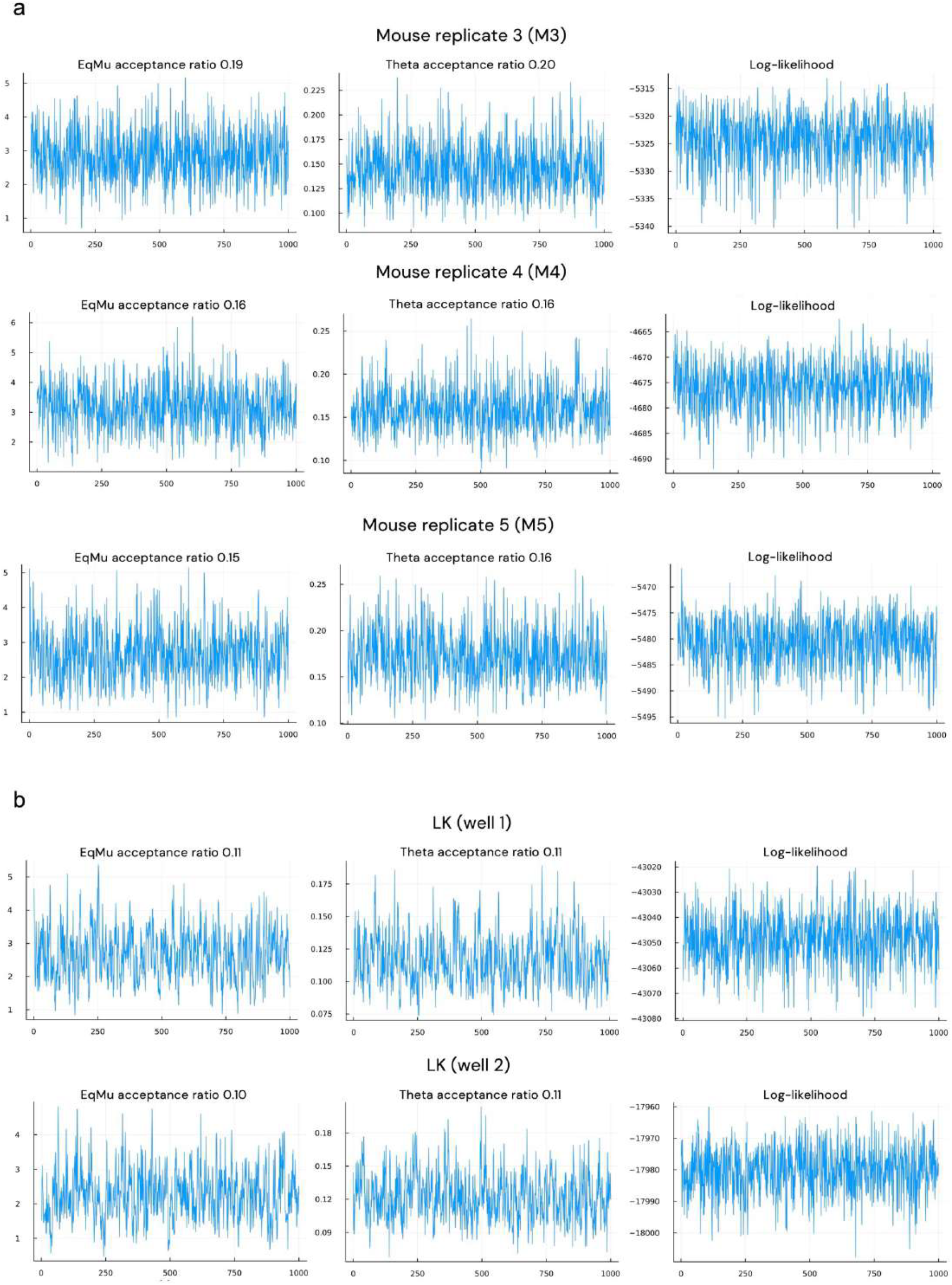
Post-burn-in MCMC trace plots for OU-model inference. Representative MCMC traces are shown for the LCMV mouse replicate analyses and the LARRY LK-subset analyses. For each dataset, traces are shown for the OU equilibrium mean (EqMu, μ; left), the mean-reversion rate (Theta, θ; middle), and the model log-likelihood (right) across the 1,000 retained post-burn-in samples. The Metropolis–Hastings acceptance ratio for each parameter proposal is annotated in the corresponding title. a. Trace plots for LCMV mouse replicates M3, M4, M5. b. Trace plots for LARRY LK wells 1 and 2.

**Figure S11.**
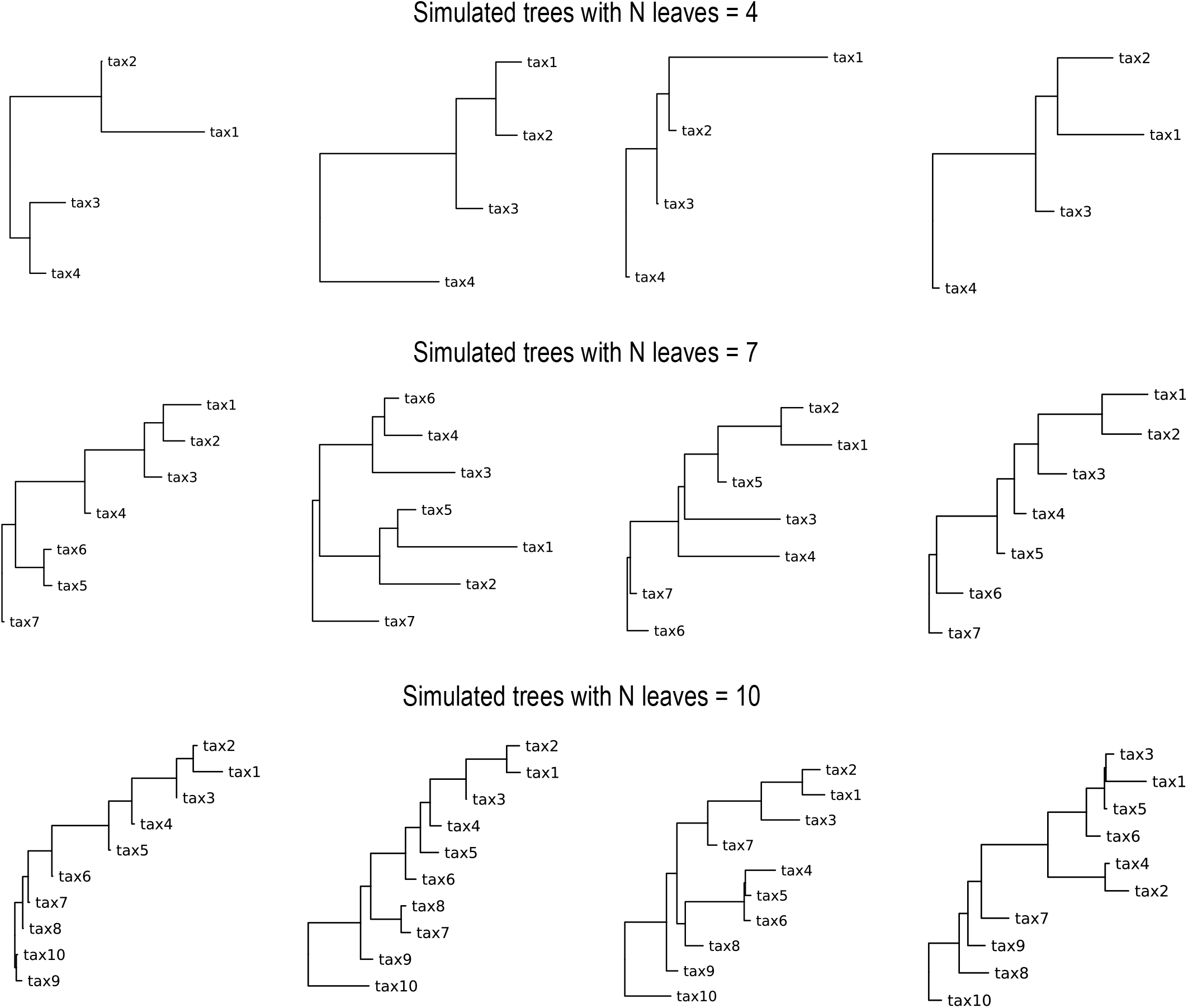
Examples of the simulated trees. Representative simulated trees used as ground truth in the simulation study, shown for trees with N = 4 (top row), N = 7 (middle row), and N = 10 (bottom row) leaves. Four random, For each tree size, multiple random, independently generated examples are displayed to illustrate the range of topological structures used in the simulations. Each simulation started from a random bifurcating tree with N leaves, and the total branch length of each tree was subsequently rescaled to 10.0 units to standardize the expected degree of clonal expansion across replicates.

**Table S1.** Simulation reconstruction accuracy Small (mean N cells = 15000)

| N Leaves | mean Pearson correlation | CI 95% lower | CI 95% upper | mean node differences | CI 95% low node differences | CI 95% high node differences |
| --- | --- | --- | --- | --- | --- | --- |
| 4 | 0.947 | 0.936 | 0.957 | 0.240 | 0.147 | 0.333 |
| 5 | 0.955 | 0.947 | 0.963 | 0.350 | 0.252 | 0.448 |
| 6 | 0.948 | 0.936 | 0.960 | 0.420 | 0.283 | 0.557 |
| 7 | 0.966 | 0.961 | 0.972 | 0.500 | 0.349 | 0.651 |
| 8 | 0.969 | 0.965 | 0.974 | 0.710 | 0.554 | 0.866 |
| 9 | 0.959 | 0.952 | 0.966 | 0.860 | 0.686 | 1.034 |
| 10 | 0.966 | 0.960 | 0.972 | 0.770 | 0.603 | 0.937 |
| Average | 0.959 |  |  | 0.550 |  |  |

Large (mean N cells = 1050000)
| N Leaves | mean Pearson correlation | CI 95% lower | CI 95% upper | mean node differences | CI 95% high node differences | CI 95% high node differences |
| --- | --- | --- | --- | --- | --- | --- |
| 4 | 0.976 | 0.966 | 0.986 | 0.070 | 0.020 | 0.120 |
| 5 | 0.992 | 0.989 | 0.994 | 0.120 | 0.050 | 0.190 |
| 6 | 0.993 | 0.991 | 0.995 | 0.150 | 0.069 | 0.231 |
| 7 | 0.992 | 0.990 | 0.994 | 0.230 | 0.134 | 0.326 |
| 8 | 0.995 | 0.994 | 0.996 | 0.210 | 0.121 | 0.299 |
| 9 | 0.989 | 0.979 | 1.000 | 0.230 | 0.103 | 0.357 |
| 10 | 0.997 | 0.996 | 0.997 | 0.220 | 0.129 | 0.311 |
| Average | 0.991 |  |  | 0.176 |  |  |

**Table S2.** Permutations tests for HDM and Dog Allergen models.

**HDM model**
| cell_type | n_cells | observed_var | null_mean | null_sd | pvalue | zscore | effect_ratio | q95 | q99 | qvalue_BH | significant_FDR_01 |
| --- | --- | --- | --- | --- | --- | --- | --- | --- | --- | --- | --- |
| IFN_response | 635 | 2663.035 | 1800.800 | 346.373 | 0.006 | 2.489 | 1.479 | 2356.933 | 2569.891 | 0.021 | FALSE |
| Th2_2 | 1356 | 277.551 | 151.291 | 59.100 | 0.036 | 2.136 | 1.835 | 259.791 | 323.174 | 0.084 | FALSE |
| Tregs | 1611 | 1350.888 | 233.713 | 75.027 | 0.001 | 14.890 | 5.780 | 373.642 | 447.143 | 0.007 | TRUE |
| Act_circ | 1099 | 1700.858 | 1694.466 | 313.901 | 0.502 | 0.020 | 1.004 | 2198.947 | 2391.487 | 0.703 | FALSE |
| Th2 | 1174 | 329.895 | 275.372 | 108.471 | 0.268 | 0.503 | 1.198 | 466.755 | 599.326 | 0.469 | FALSE |
| Naive | 319 | 97.745 | 113.278 | 23.268 | 0.744 | -0.668 | 0.863 | 151.240 | 167.415 | 0.744 | FALSE |
| Proliferating | 99 | 168.102 | 538.974 | 571.982 | 0.606 | -0.648 | 0.312 | 1600.904 | 2300.785 | 0.707 | FALSE |

**HDM model: reclustered Tregs**
| cell_type | n_cells | observed_var | null_mean | null_sd | pvalue | zscore | effect_ratio | q95 | q99 | qvalue_BH | significant_FDR_01 |
| --- | --- | --- | --- | --- | --- | --- | --- | --- | --- | --- | --- |
| Tregs-0 | 474 | 41.238 | 87.090 | 82.559 | 0.656 | -0.555 | 0.474 | 235.195 | 390.561 | 0.737 | FALSE |
| Tregs-1 | 462 | 25.161 | 15.703 | 22.516 | 0.191 | 0.420 | 1.602 | 59.808 | 104.074 | 0.649 | FALSE |
| Tregs-2 | 370 | 3460.295 | 3000.278 | 804.829 | 0.325 | 0.572 | 1.153 | 4132.702 | 4425.725 | 0.649 | FALSE |
| Tregs-3 | 305 | 417.588 | 593.332 | 254.656 | 0.737 | -0.690 | 0.704 | 1048.437 | 1316.138 | 0.737 | FALSE |

**Dog Allergen model**
| cell_type | n_cells | observed_var | null_mean | null_sd | pvalue | zscore | effect_ratio | q95 | q99 | qvalue_BH | significant_FDR_01 |
| --- | --- | --- | --- | --- | --- | --- | --- | --- | --- | --- | --- |
| Tfh | 566 | 3514.026 | 3529.861 | 493.467 | 0.542 | -0.032 | 0.996 | 4313.102 | 4607.558 | 0.678 | FALSE |
| Naive | 354 | 0.330 | 0.569 | 0.113 | 0.990 | -2.108 | 0.580 | 0.758 | 0.814 | 1.000 | FALSE |
| Act_circ | 597 | 1344.938 | 727.650 | 282.271 | 0.022 | 2.187 | 1.848 | 1243.239 | 1577.873 | 0.155 | FALSE |
| Th2 | 884 | 296.362 | 182.273 | 78.622 | 0.086 | 1.451 | 1.626 | 328.995 | 396.955 | 0.265 | FALSE |
| Tregs | 375 | 154.132 | 105.775 | 84.374 | 0.222 | 0.573 | 1.457 | 270.703 | 389.834 | 0.444 | FALSE |
| Th_CTL_like | 220 | 6490.070 | 5939.309 | 1180.163 | 0.330 | 0.467 | 1.093 | 7809.101 | 8467.768 | 0.471 | FALSE |
| Act2 | 759 | 612.445 | 304.740 | 145.544 | 0.031 | 2.114 | 2.010 | 577.789 | 758.671 | 0.155 | FALSE |
| Act3 | 262 | 313.117 | 144.370 | 130.988 | 0.106 | 1.288 | 2.169 | 397.370 | 614.981 | 0.265 | FALSE |
| Th17 | 1012 | 727.934 | 611.441 | 209.437 | 0.289 | 0.556 | 1.191 | 994.743 | 1117.000 | 0.471 | FALSE |
| IFN_response | 51 | 0.000 | 6.464 | 28.191 | 1.000 | -0.229 | 0.000 | 6.464 | 129.283 | 1.000 | FALSE |

**Table S3.**
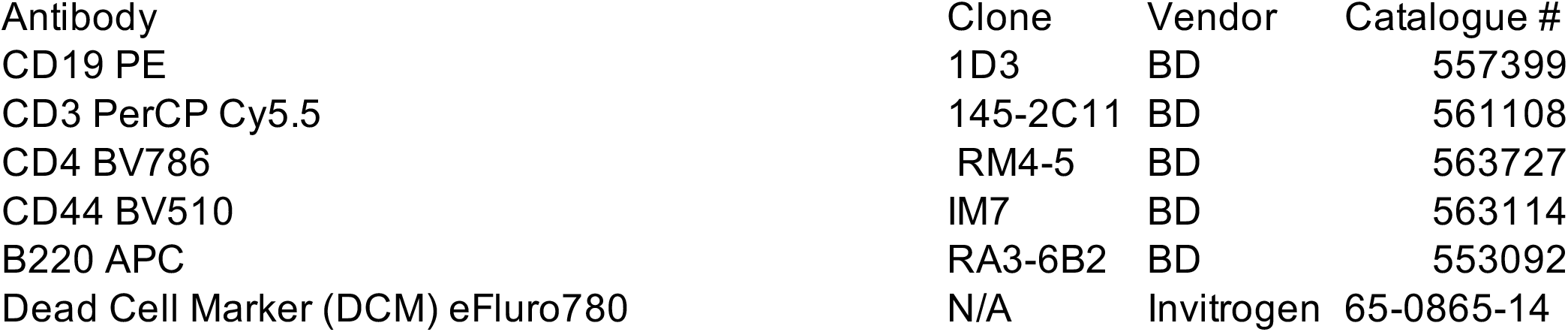
Antibodies used for single-cell staining.

| Antibody | Clone | Vendor | Catalogue # |
| --- | --- | --- | --- |
| CD19 PE | 1D3 | BD | 557399 |
| CD3 PerCP Cy5.5 | 145-2C11 | BD | 561108 |
| CD4 BV786 | RM4-5 | BD | 563727 |
| CD44 BV510 | IM7 | BD | 563114 |
| B220 APC | RA3-6B2 | BD | 553092 |
| Dead Cell Marker (DCM) eFluro780 | N/A | Invitrogen | 65-0865-14 |

